# WAChRs are excitatory opsins sensitive to indoor lighting

**DOI:** 10.1101/2025.09.12.675947

**Authors:** Amanda J. Tose, Alberto A. Nava, Sara N. McGrath, Alan R. Mardinly, Alexander Naka

## Abstract

Hundreds of novel opsins have been characterized since the advent of optogenetics, but low experimental throughput has limited the scale of opsin engineering campaigns. We modified an automated patch-clamp system with a multispectral light source and a custom light path to enable high-throughput electrophysiological measurements of opsin functional properties. Using this approach, we screened over 1,750 opsins from a range of families. We discovered that the F240A mutation of the light-gated potassium channel WiChR abolished potassium selectivity, turning it into a sensitive excitatory channel that we dubbed “WAChR”. We systematically mutated WAChR and identified variants that expand the frontier of speed-sensitivity tradeoffs. Multiple WAChR variants produced large inward currents in response to indoor ambient office light, and responded to irradiances as low as 15 nW/mm^2^, something that we did not observe with other ultra-sensitive opsins. *In vivo* recording from the mouse cortex confirmed that WAChRs exhibit enhanced sensitivity in neurons. These ambient-light sensitive channels should be broadly useful for neuroscience research and vision restoration therapies.

## Introduction

The use of channelrhodopsin (CR) variants for optogenetics has revolutionized neuroscience by allowing neurons to be activated by light upon expression of a light-gated ion channel. From the earliest days of optogenetics, scientists have understood the therapeutic potential of CRs for vision restoration in patients with photoreceptor degeneration^1^. CRs are particularly well-suited for vision restoration due to their millisecond-scale kinetics and single-component simplicity. However, since CRs lack the biochemical amplification of native photoreceptors, they are orders of magnitude less sensitive to light than endogenous phototransduction^2,3^. This has required optogenetic clinical trials to equip patients with light-amplifying goggles that project intense light onto the retina^4^. In addition to safety concerns, these devices impose practical barriers, such as limited field of view and eye-box, social stigma, and substantial power requirements. The development of CRs sensitive enough to function under natural indoor lighting conditions without assistive devices would therefore be transformative for clinical applications of optogenetics.

To address this challenge, the field has pursued both discovery and engineering approaches over the past two decades. Systematic screening of microbial genomes has dramatically expanded the CR toolkit, revealing diverse natural variants with distinct spectral, kinetic, and conductance properties^5–12^. In parallel, researchers have employed rational design and computational approaches to engineer improved CRs, creating mutants and chimeric proteins with altered spectral tuning, modified kinetics, enhanced conductance, changes in ion selectivity, improved expression and membrane trafficking, and in some cases, better light sensitivity^13–49^. These engineering efforts have leveraged structural insights, directed evolution, and increasingly sophisticated screening methods to push the boundaries of CR performance. Most recently, the discovery of pump-like channelrhodopsins (PLCRs) has been particularly significant, yielding both highly sensitive cation-conducting variants like ChRmine and the first potassium-selective channelrhodopsins (KCRs), which combine novel ion selectivity with exceptional baseline sensitivity^6,7,40,41,50,51^.

Yet despite these advances, the sensitivity of available CRs, especially for low light applications such as vision restoration, still leaves much to be desired^3,52,53^. One explanation for this is a fundamental technical bottleneck: manual whole-cell patch-clamp electrophysiology, the gold standard for characterization, is slow and labor-intensive. This low throughput makes it exceptionally difficult to navigate the complex fitness landscape of CRs, where improvements in one property, like light sensitivity, can come at the cost of another, such as kinetics or expression. Without the throughput to systematically explore vast sequence spaces or generate the large functional datasets needed for machine learning approaches, the field remains unable to engineer the transformative variants required for practical clinical applications.

To overcome these limitations, we developed a high throughput assay based on a modified microfluidic patch-clamp system. We deployed this assay as a core component of an active learning loop, leveraging protein language models to explore the CR fitness landscape and iteratively optimize promising variants. Early in our screen, we discovered the F240A mutation in the KCR WiChR, which transforms it into a potent nonspecific cation channel. We dubbed this exceptionally sensitive excitatory channel “WAChR”. Our campaign identified a suite of WAChR variants that are situated along the frontier of speed versus sensitivity, allowing for tradeoffs between these properties. Notably, many WAChRs produced large photocurrents in response to standard indoor office lighting large enough to evoke action potentials in neurons. Unlike other opsins advanced as candidates for vision restoration, WAChRs exhibited photocurrent responses to nanowatt irradiances. We further validated the exceptional light sensitivity and biocompatibility of WAChRs in mouse brain using viral vectors. This new suite of ultra-sensitive opsins should enable ambient light optogenetic experiments and therapeutics.

## Results

### A high throughput assay of channelrhodopsin functional properties

The usefulness of optogenetic proteins is a combination of numerous functional properties, including light sensitivity, spectral selectivity, kinetics, and macroscopic photocurrent amplitude. To systematically engineer these properties, we developed an approach that evaluates them all in the same experiment while retaining the throughput needed for a large opsin engineering campaign. To accomplish this, we modified a commercially available plate-based automated patch-clamp system (the Ionflux Mercury 16; Cell Microsystems) to accommodate an optical stimulation path (Fig 1A, Fig S1A). We used a bank of 7 different solid-state light sources to generate light at a range of wavelengths and irradiances, and used a galvo-galvo system to steer these photostimuli onto the different recording sites on the plates (Fig S1B-D). We transfected opsins of interest in HEK293T cells using a standardized expression cassette, obtained whole-cell patch recordings, and made measurements enabling us to estimate the sensitivity, spectral selectivity, photocurrent amplitude, and kinetics on each recording. To maximize the likelihood of success in a given recording, we used “ensemble” plates, which are configured such that each electrode records from 20 micropipettes in parallel (Fig 1B-C, Fig S1E-F).

**Figure 1.**
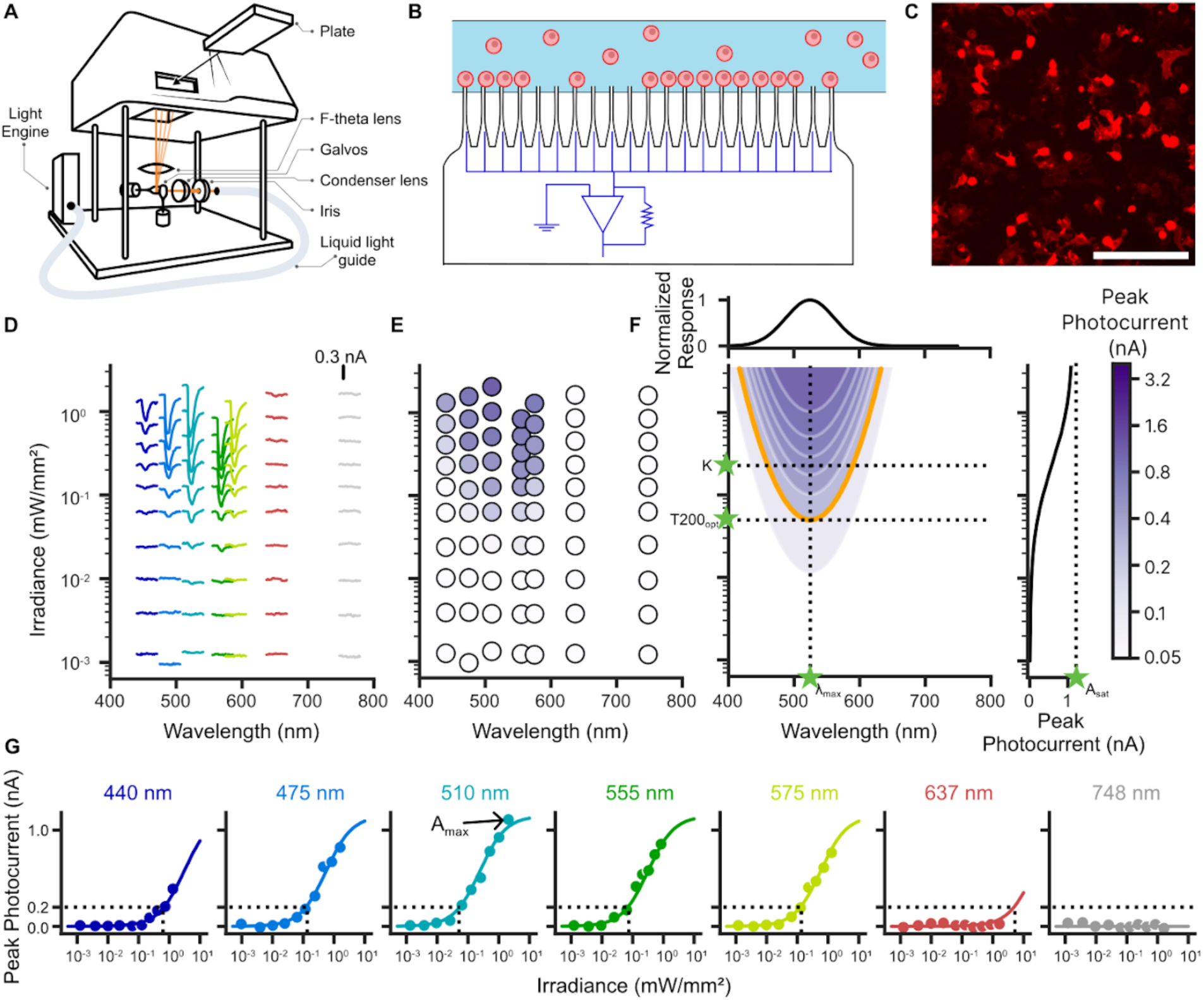
A platform for high-throughput channelrhodopsin screening (A) Schematic of the automated patch-clamp system and light path. (B) Illustration of an individual recording site where 20 micropipettes are connected in parallel to perform an ensemble electrophysiology recording of up to 20 cells. CR-expressing cells (red) flow through microfluidic channels (light blue) to reach the recording site where there are 20 micropipettes (black) configured to record in parallel (dark blue). (C) Representative fluorescence microscopy image of ChRmine-expressing HEK239T cells. All opsins in our screen were expressed with a C-terminal mScarlet tag enabling visualization. Scale bar is 200 µm. (D) Representative median photocurrent traces produced from a single channel of an automated patch-clamp experiment for ChRmine for different photostimulation conditions. Traces are positioned on the x and y axes according to the wavelength and irradiance of the corresponding photostimulus. Each photostimulus was 90ms in duration; relative to stimulus start time, traces cover from -100 ms until +290 ms (390 ms total duration for each trace). (E) Peak photocurrent amplitudes are measured from each trace in (D). (F) A response surface is fit to enable a view of opsin photocharacteristics. The orange contour line represents the irradiance required to evoke 200 pA at any given wavelength. The top panel shows the normalized spectral selectivity while the right panel shows the irradiance dose response function. Breakdown of responses by wavelength. Scatterplots show the median photocurrent amplitude versus irradiance for each of 7 different wavelengths; each scatterplot is essentially a vertical slice through one of the wavelengths portrayed in (E). Curves show the conditional predictions of the fitted action surface model for a range of irradiances at the corresponding wavelength; these are essentially vertical slices through the contour map portrayed in (F). Dashed black lines on each subplot indicate the 200 pA photocurrent threshold and the irradiance required to evoke 200 pA at that wavelength.

To establish experimental parameters for our campaign, we ran pilot experiments using the pump-like channelrhodopsin ChRmine^7,40,54^ as our baseline opsin. To mitigate trafficking, folding, and expression-related failures, we incorporated motifs which have been reported to improve expression and membrane trafficking efficiency^17,28,46,47,55–57^. These include the Kir2.1 Golgi trafficking sequence (G), a fluorophore (F; typically mScarlet), the soma-targeting sequence from Kv2.1 (S), and the Kir2.1 endoplasmic reticulum export sequence (E). This collection of sequences was fused to the C-terminus of the opsin in the order GFSE (Fig S1G-H). In some cases, we also prepended the cleavable LucyRho (LR) tag to the N-terminus of the opsin as an additional measure to enhance surface expression^58–60^. We optimized transfection conditions and promoters and decided to employ the strong constitutive promoter EF1a to drive opsin expression in HEK293T cells (Fig S2).

We next designed our light stimulus conditions. Established approaches to characterizing CR function include measuring photoresponses at a fixed wavelength while varying the intensity of photostimulation (the dose response function) and measuring photoresponses at varying wavelengths while fixing or controlling for photon flux (the action spectrum). These sets of measurements are typically performed as separate experiments, but we opted to probe both dimensions of photostimulation simultaneously. We developed a protocol in which we applied 90 ms photostimuli at 7 different wavelengths (spanning 440 to 748 nm) and at 10 different irradiances for each wavelength (∼1 µW/mm² to 1 mW/mm²) while recording photocurrent responses (Fig 1D-E). This allowed us to visualize the relationship between wavelength, irradiance, and photocurrent, which we term the “action surface” (Fig 1F).

We developed a simple two-stage model to describe the observed action surface of a single CR. First, a spectral efficiency filter (essentially the action spectrum; approximated by a Gaussian) scales the incoming irradiance (E) based on wavelength (λ) to determine an effective irradiance (E_λ_). This effective irradiance is then used as the input to the Hill equation to model the dose response relationship (Fig 1G). This model yielded a set of parameters that concisely captured the functional properties of each opsin including its peak activation wavelength (λ_max_), saturated response amplitude (A_sat_), and half-saturation constant (K; sometimes referred to as EC50). For recordings where the median photocurrent exceeded 200 pA for at least one of the stimulation conditions, we typically observed good fits to the overall action surface (median R^2^ = 0.96, Fig S3A-B). Ideally, the A_sat_ parameter would correspond to the photocurrent that would be observed in a given recording if the opsin was stimulated with saturating levels of light, but we did not achieve irradiances this high for most opsins in our primary assay. To complement this, we also calculate a model-free metric, A_max_, which is simply the value of the median photocurrent amplitude for the stimulation condition with the largest response; in practice this was very closely correlated with A_sat_.

For a CR to be effective at driving spiking in neurons via low intensity photostimulation, it needs to be both sensitive and capable of generating large magnitude photocurrents. As an integrative measure, we used the conditional predictions of the model to estimate the threshold irradiance required to evoke 200 pA of photocurrent. As this value is dependent on the wavelength of the photostimulation, we calculated it for each individual wavelength probed in our assay (e.g. we use T200_475_ to denote the 200 pA threshold irradiance stimulating at 475 nm) as well as for the predicted optimal wavelength (T200_opt_). This metric is both reliable (since it is essentially an interpolation) and interpretable in practical terms; we therefore used it as our primary measure of efficacy. We also calculated the decay time constant of photoresponses (τ_off_).

### Creating a baseline dataset

One of the challenges in performing sequence-based predictions of CR functional properties is that experimental and analytical methods vary widely in existing datasets. Having established our assay conditions and parameters, we sought to obtain baseline measurements from a wide range of published optogenetic tools to generate a standardized dataset and identify promising scaffolds for further engineering. Here, we focus mainly on excitatory cation-selective channelrhodopsins (CCRs) and pump-like channelrhodopsins (PLCRs). Consistent with what has previously been reported, we observed low T200_opt_ values for variants of green algal CCRs specifically engineered for enhanced sensitivity, such as ex3mV1Co^34^, ChRger2^15^, and CoChR-3M^33^ (Fig 2A-B). Many of the recently identified PLCRs have also been reported to be exceptionally sensitive. Consistent with this, HcCCR^61^, GtCCR4^13,62^, ChRmine, and the engineered ChRmine variant ChReef^45^ ranked among the most sensitive CRs (Fig 2A-B). The PLCR family also includes the KCRs HcKCR1^6^, HcKCR2, and WiChR^50^. We observed modest currents from wildtype KCRs, likely due to the holding potential in our assay being close to the reversal potential for potassium (Fig 2B). However, we did see sizeable responses from HcKCR1/2 variants with the Y222A mutation^41,50^, which converts them from K^+^ selective channels into non-selective cation channels (Fig 2B). We applied the analogous F240A mutation to WiChR, which is reported to be more sensitive than the HcKCRs^50^. We saw promising results: relative to other CRs, WiChR F240A produced large photocurrents while maintaining excellent sensitivity (Fig 2A-B, WiChR F240A: T200_opt_ = 3.1 µW/mm², K = 18 µW/mm², A_max_ = 3.9 nA, τ_o_ = 260 ms, λ_max_ = 484 nm, n = 2; ChRmine T200_opt_ = 86.5 µW/mm², K = 232 µW/mm², A_max_ = 0.52 nA, τ_o_ = 79.6 ms, λ_max_ = 523 nm, n = 39).

**Figure 2.**
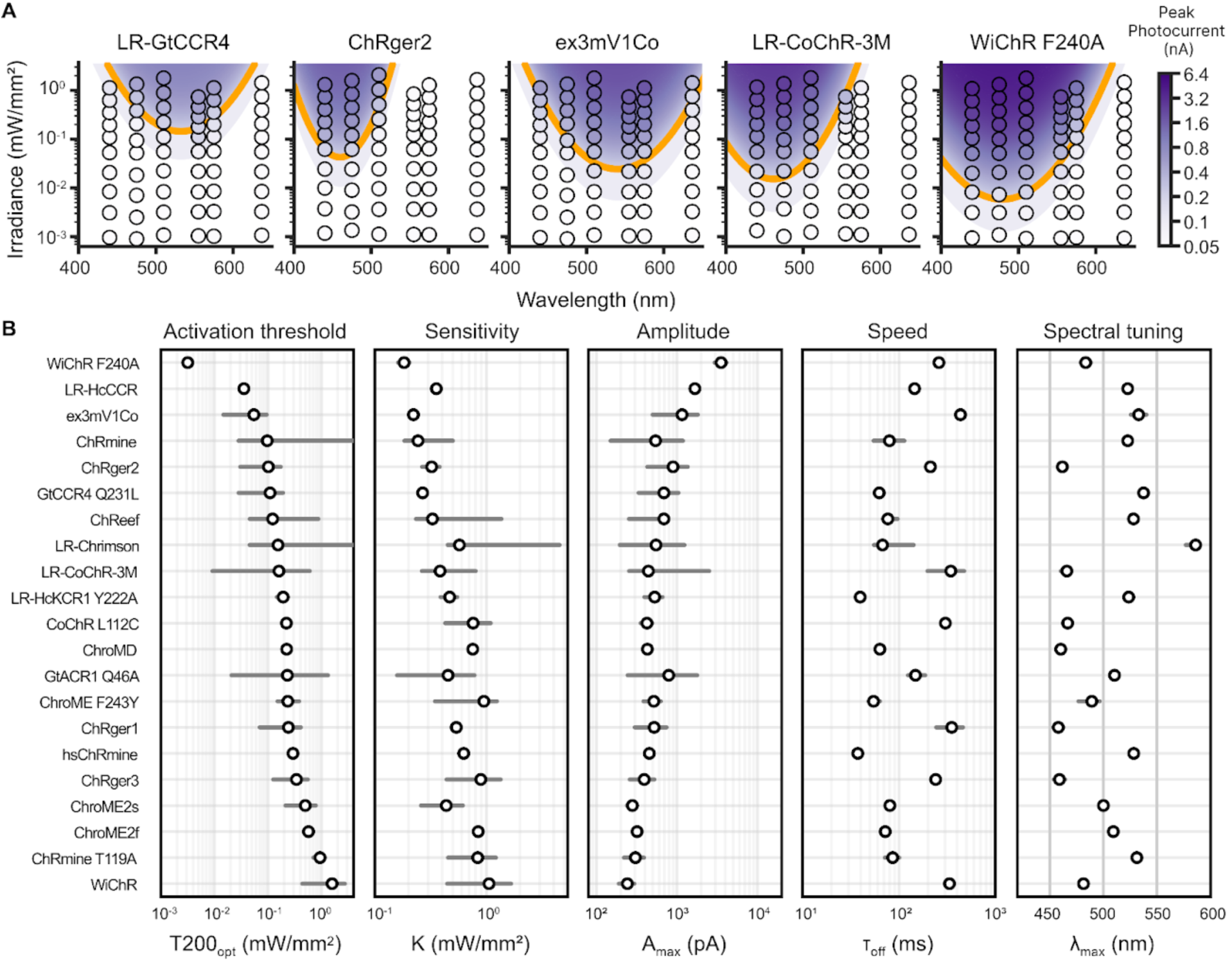
A preliminary screen of known opsins identifies WiChR F240A as a leading candidate for high-sensitivity applications (A) Representative action surfaces comparing the novel WiChR F240A mutant to other well-characterized sensitive opsins. Each plot displays the peak photocurrent (color bar) as a function of light wavelength and irradiance. The orange contour line represents the modeled threshold irradiance required to evoke 200 pA of current (T200), a primary measure of efficacy. Quantitative comparison of five key performance metrics across a broader library of opsins. The metrics evaluated are: Activation threshold (T200_opt_): The irradiance at the optimal wavelength needed to elicit a 200 pA response. Sensitivity (K): The half-saturation constant, where lower values indicate higher sensitivity. Amplitude (A_max_): The maximum recorded photocurrent. Speed (τ_off_): The deactivation time constant after light cessation. Spectral tuning (λ_max_): The wavelength of peak activation. Data from the median (white dots with black outline) and range (gray bars) are plotted for each opsin.

### WAChR is an excitatory optogenetic tool

Structural and mutational analyses of HcKCR1 have shown that the residues N99, W102, and Y222 (N117, W120, and F240 in WiChR) form the K^+^ selectivity filter with support from nearby residue C29 (D47 in WiChR)^41,50,63^ (Fig 3A). Because mutations to these residues can disrupt K^+^ selectivity, we hypothesized that the F240A mutation in WiChR would have a similar effect, analogous to the Y222A mutation in HcKCR1 (Fig 3B, Fig S4). Using a manual patch-clamp electrophysiology rig for improved data quality, we performed a current-voltage (I-V) analysis for WiChR and WiChR F240A. This experiment showed that the reversal potential for WiChR F240A is roughly 75 mV more positive than wildtype WiChR under standard recording conditions, likely due to increased sodium permeability (Fig 3C-D, WiChR F240A: -6.8 ± 1.1 mV vs. WiChR: -81.8 ± 2.7 mV, Mann-Whitney U test, p < 0.001, n = 21 and 10 respectively). By shifting the photocurrent reversal potential above the typical action potential threshold, the F240A mutation effectively converts WiChR into an excitatory optogenetic tool, while retaining the large photoconductance, high sensitivity, and minimal desensitization of wildtype WiChR^50^. We named this new variant “WAChR”.

**Figure 3.**
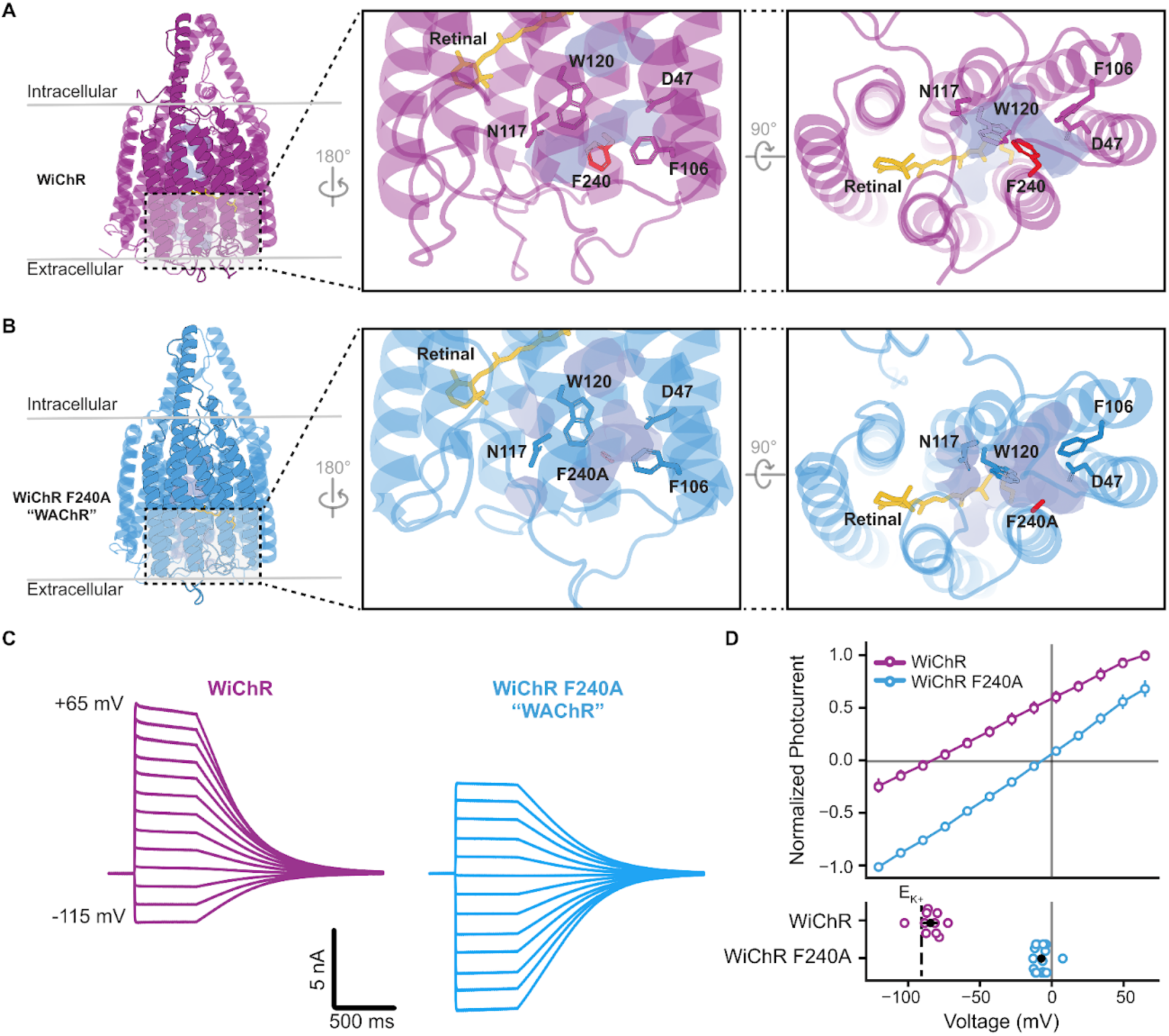
The F240A mutation abolishes K⁺ selectivity, converting WiChR into an excitatory channel (WAChR) (A, B) Predicted protein structures of the wild-type WiChR (A) and the F240A mutant, WAChR (B). The expanded views on the right highlight the K⁺ selectivity filter. In WAChR, the bulky phenylalanine at position 240 is replaced with a smaller alanine residue (F240A), disrupting the filter. (B) Representative photocurrent traces from manual patch-clamp recordings of WiChR (purple) and WAChR (blue) in response to 500 ms of 480 nm light, recorded at various holding voltages ranging from -115 mV to +65 mV. The current-voltage (I-V) relationship (top) and summary data (bottom) show a significant positive shift in the reversal potential for WAChR compared to WiChR. This shift confirms that the F240A mutation converts the channel from potassium-selective (inhibitory) to non-selective cation-permeable (excitatory).

### A campaign to optimize WAChR

Having identified WAChR as the most efficacious excitatory CR in our preliminary screen, we directed our machine learning guided engineering campaign towards improving the functional properties of WAChR. We refined our methods throughout the campaign, but broadly followed established frameworks^64^. Following earlier foundational work^15,16^, we compiled publicly available information and combined this with our preliminary data to assemble a dataset of functionally labeled CR sequences. We used this dataset to train ensembles of sequence-to-function models based on fine-tuned pretrained protein language models. We generated candidates through a variety of means (see Methods) that were ultimately filtered and ranked by our models. We tested sequences in batches and used the resulting data to enrich our dataset for the next design cycle. Overall, we tested 672 unique WAChR opsin variants from 13 batches of synthesis and 2 batches of mutagenesis. We observed changes in sensitivity, photocurrent magnitude, and kinetics (Fig 4A,B, Fig S5). Notably, most of the mutations that we made to WAChR did not eliminate channel activity (212 of 672 produced no measurable photocurrent), but most mutations did impair the efficacy, sensitivity, and peak photocurrent amplitude (Fig 4A, Fig S5).

**Figure 4.**
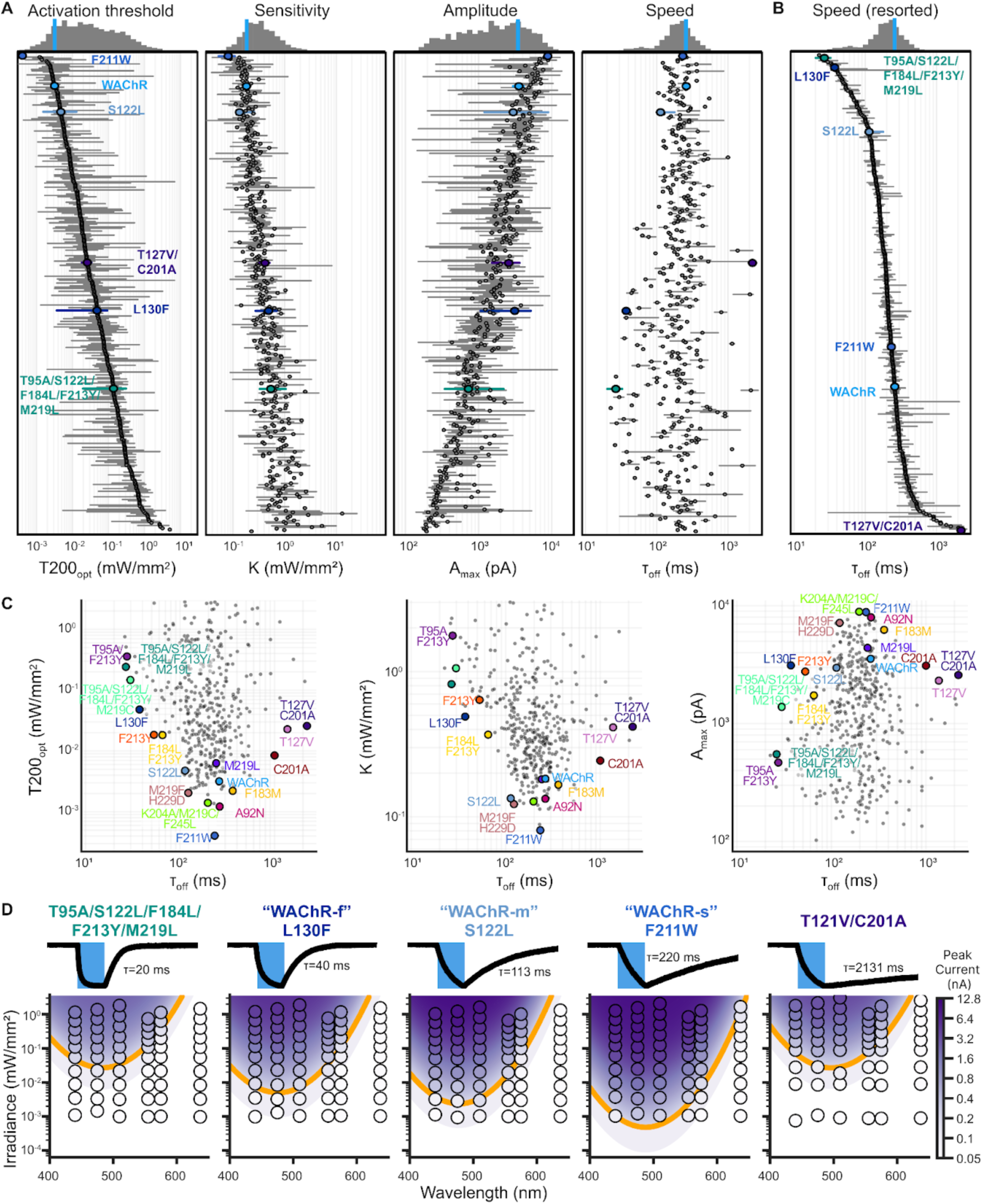
A high-throughput engineering campaign expands the speed-sensitivity frontier of WAChR variants (A) Distribution of key functional properties measured across 672 unique WAChR variants. The plots show the activation threshold (T200_opt_), sensitivity (K), photocurrent amplitude (A_max_), and speed (τ_off_). Individual variants are shown in gray, with several notable mutants highlighted in color. The histograms at the top of each plot summarize the distribution for each metric. The blue bar indicates the position of WAChR in the distribution. (B) The speed (τ_off_) data from panel A, resorted to show the ranking of variants from fastest (top) to slowest (bottom). (C) Scatter plots illustrate the trade-offs between off-kinetics (τ_off_) and the other primary performance metrics: efficacy (T200_opt_), sensitivity (K), and amplitude (A_max_). This analysis reveals a frontier of variants that achieve optimal combinations of speed and sensitivity. Key variants residing on or near this frontier are labeled. Representative photocurrent traces (top) and corresponding action surfaces (bottom) for five selected variants that span a wide range of kinetics. The traces are annotated with their deactivation time constant (τ_off_).

Changes to CR kinetics often involve a tradeoff with sensitivity: accelerating kinetics typically reduces sensitivity, while decelerating kinetics can increase sensitivity^13,23,32,49,65^.

However, some tradeoffs are better than others: for example, not every mutation that slows kinetics also increases sensitivity^13,32^. We identified a number of individual mutations that accelerate the off-kinetics of WAChR, including S122L, L130F, F184L, F213Y, and F220Y, as well as mutations that decelerated them, including T127V, F183M, and C201A (Fig 4B-D).

Some of these sites are located on the interface between the 5th transmembrane (TM) domain and adjacent TM domains, where kinetic-altering mutations have previously been identified^66^; for example, the F213 and F220 residues of WAChR are likely homologous with Y261 and Y268 in Chrimson.

Notably, some of these variants, such as S122L, M219F/H229D, and K204A/M219C/F245L, accelerated the kinetics of WAChR without dramatically reducing the sensitivity (Fig 4C). We therefore turned our attention toward identifying sets of mutations that would yield favorable tradeoffs between off-kinetics and sensitivity. As the campaign progressed, we expanded the frontier in the domain of τ_off_ and T200_opt_ (Fig 4C). We selected three high-performing variants along this frontier as candidate tools: L130F, S122L, and F211W. For simplicity we refer to these variants as WAChR-f, WAChR-m, and WAChR-s (fast, medium and slow, Fig 4D).

### WAChRs are ultrasensitive

To better characterize the response properties of our frontier WAChR variants, we returned to manual patch-clamp recordings. Since most optogenetic applications depend on viral delivery, we synthesized adeno-associated viruses (AAVs) of our leading WAChR variants, expressing it under the ubiquitous EF1a promoter. We noticed that morphologically, AAV-transduced WAChR HEK293T cells appeared healthier than cells lipofectamine transfected with the same WAChR variant (Fig S6A). To more fairly benchmark WAChRs against other state of the art opsins, we packaged AAVs encoding CoChR-3M, ex3mV1Co, ChroME2s, and ChReef ^44,67,7,40,45,32,33,68,34^. To emulate prior work examining ex3mV1Co and CoChR-3M for vision restoration applications, we used constructs in which these opsins were fused only to a fluorophore (mScarlet), without any signal peptides^33,34^. All other opsins were fused to our standard GFSE suffix. Directly comparing estimates of opsin properties from AAV or lipofection-mediated DNA delivery via manual patch revealed that many of the frontier opsins performed significantly better in the context of AAV transduction (Fig S6A-B). Notably, this was not true of WAChR-f/m/s, suggesting that their excellent performance in our screen could be partially attributed to a propensity to express and traffic well via lipofection (Fig S6B). For the remainder of the manuscript we compare opsins solely using AAV-mediated gene delivery.

To benchmark WAChR variants against other frontier opsins using AAVs, we obtained manual patch-clamp recordings and used a tunable light source to stimulate each opsin at approximately its maximally preferred wavelength at irradiances spanning several orders of magnitude (Fig 5A, Fig S7A-B, WAChR-s: n=10, WAChR-m: n=19, WAChR-f: n=11, ChReef: n=17, CoChR-3M: n=17, ex3mV1co: n=15, ChRmine: n=10, ChroME2s: n=10 cells). As with the automated patch-clamp assays, we measured T200_opt_, K, A_max_, and τ_off_ (Fig 5B). The WAChR variants all produced significantly larger maximal photocurrents (A_max_) than ChReef,

**Figure 5.**
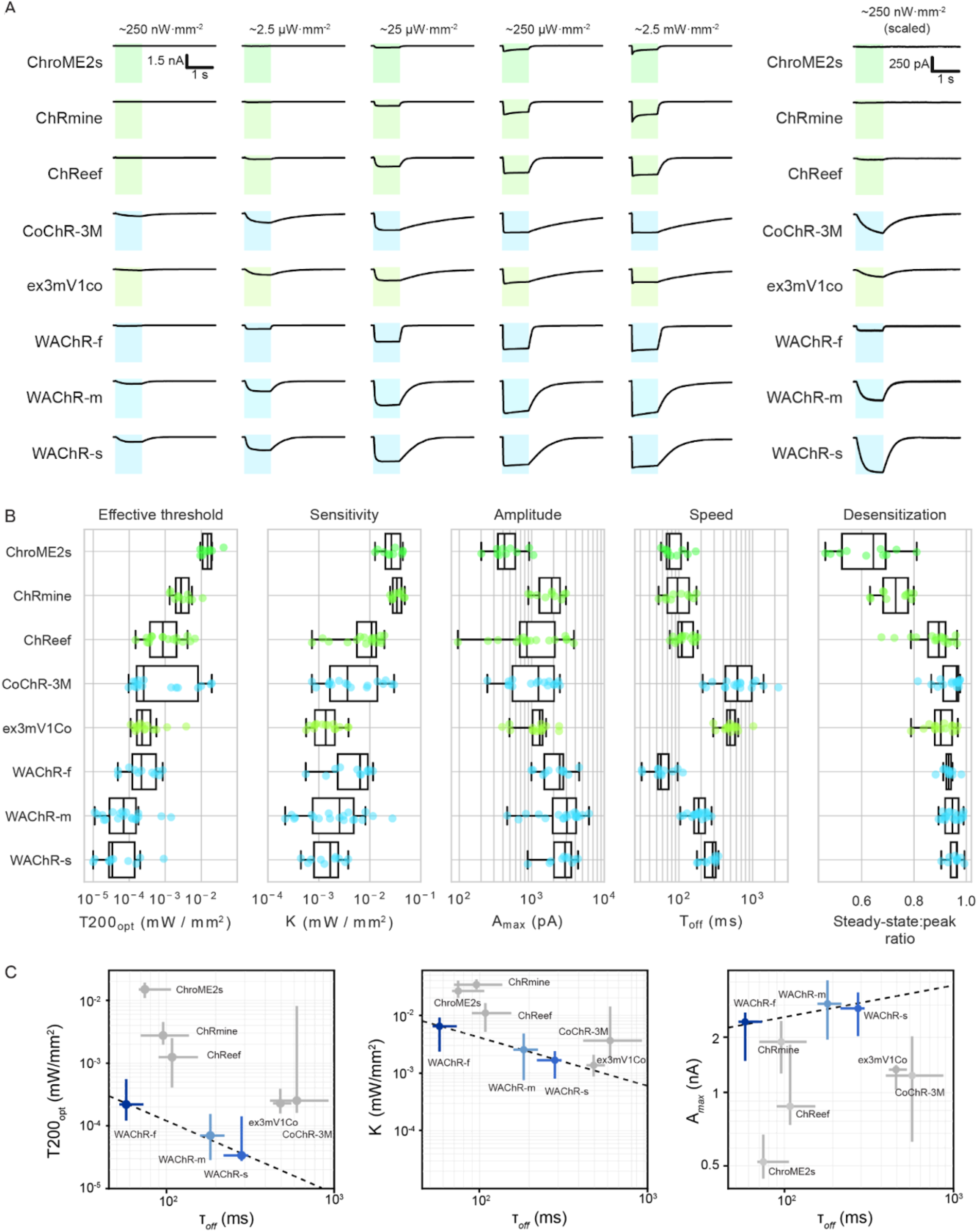
Detailed sensitivity benchmarks of AAV-transduced opsins (A) Representative photocurrent traces obtained via manual patch-clamp recordings under 1 second illumination using the λ_max_ wavelength for each opsin. Traces in the rightmost column are re-scaled versions of the traces in the first column. The wavelength of light is indicated by the color. Exact irradiances varied between experiments because of the spectrally non-uniform power output of the arc lamp in the tunable light source (see Methods). (B) Distributions of photocurrent metrics obtained from each opsin. Each dot indicates a recording from one cell, colored based on stimulation wavelength. Metrics of efficacy (T200_opt_), sensitivity (K), and amplitude (A_max_) are plotted versus kinetics (τ_off_) for each opsin. Dashed lines indicate the current frontier of speed/efficacy tradeoffs.

CoChR-3M, and ex3mV1Co (all p<0.05, FDR-corrected Welch’s t-test). Comparisons of the half-saturation constant (K), presented a more nuanced picture. WAChR-s was significantly more sensitive than both ChReef (p<0.001) and CoChR-3M (p<0.05). WAChR-m was more sensitive than ChReef (p<0.05) but not significantly different from CoChR-3M. When compared to ex3mV1Co, neither WAChR-s nor WAChR-m showed a significant difference in sensitivity (p>0.05), while WAChR-f was somewhat less sensitive (p<0.05). However, both WAChR-s and WAChR-m required significantly less light to evoke a 200 pA current (T200_opt_) than all three benchmark opsins (all p<0.01). WAChR-f also showed a significantly lower activation threshold than ChReef (p<0.01) and CoChR-3M (p<0.05). Kinetically, the WAChR variants offered significant speed advantages over other highly sensitive opsins. All three WAChR variants were significantly faster than both CoChR-3M and ex3mV1Co (all p<0.001). Furthermore, WAChR-f was also significantly faster than the fast benchmark ChReef (p<0.001, Fig 5B). Taken together, these results establish WAChRs as frontier tools within this group of highly sensitive opsins (Fig 5C). WAChR-s and WAChR-m offer superior T200_opt_, A_max_, and τ_off_ compared to CoChR-3M and ex3mV1Co. WAChR-f shows a similar edge when compared to the faster opsins ChReef, ChRmine, and ChroME2s. Of note, WAChR-m’s sensitivity did not depend upon fusion to the fluorophore, as a version in which the fluorophore was cleaved via a P2A site was similarly effective (Fig S6C).

The majority of these experiments were conducted using a broad-band configuration on our tunable light source which allowed stimulation at irradiances ranging from ∼250 nW/mm^2^ up to ∼2.5 mW/mm^2^ (Fig 5A, Fig S7A,B). Since we observed strong responses for some opsins even at the low end of that range, we performed an additional set of measurements on the most sensitive opsins using a narrow-band configuration which could photostimulate at irradiances on the order of 10 nW/mm^2^ (Fig S7C, Fig S8). WAChR-s, WAChR-m, and WAChR-f all exhibited reliable and reproducible responses to irradiances of ∼15 nW/mm^2^ (Fig 6A) or even lower (Fig S7C, Fig S8). Conversely, ex3mV1Co, CoChR-3M, and ChReef did not produce discernable responses at these irradiances (Fig 6B) and only started to yield photocurrents at irradiances of at least 30 nW/mm^2^ (Fig S7C). The evoked responses of WAChR-s (95.1±49.2 pA, n = 9 cells) and WAChR-m (46.4±28.1 pA, n = 6 cells) were each significantly larger than those of the benchmark opsins (ex3mV1Co: 2.9±5.5 pA, n = 7 cells; CoChR-3M: 4.1±2.2 pA, n = 4 cells; ChReef: -0.9±1.5 pA, n = 6 cells; p<0.05, all comparisons p<0.05, FDR-corrected Welch’s t-test ) and the response of WAChR-f (30.0±18.4 pA, n = 8 cells) was significantly larger than that of ChReef (p<0.05) but not significantly different than that of CoChR-3M (p=0.052) or ex3mV1Co (p=0.051).

**Figure 6.**
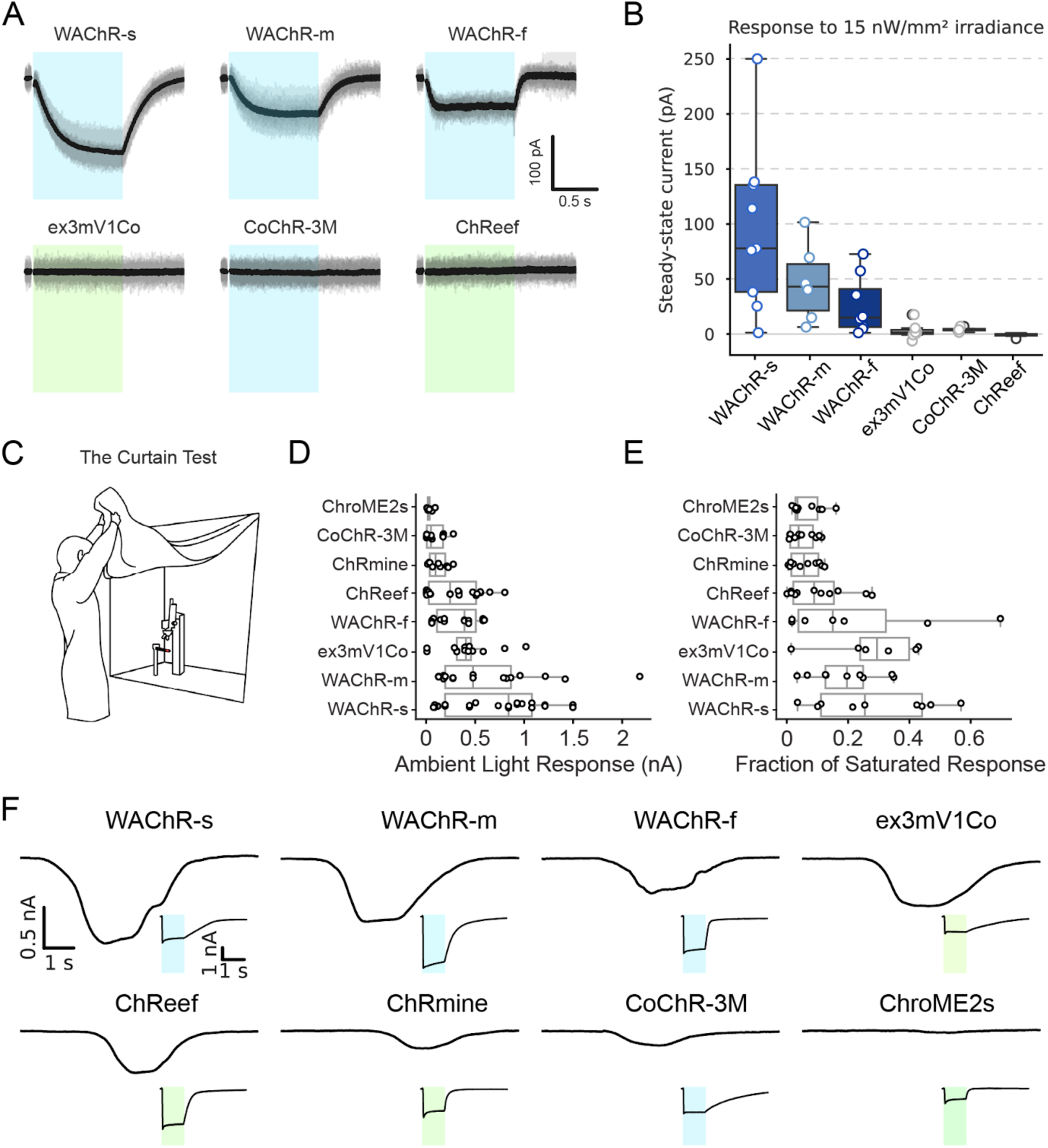
WAChR variants are ambient light sensitive (A) Representative photocurrent traces showing responses to illumination at irradiances of 9-13 nW/mm². The example trace for ChReef is at an irradiance of approximately 9 nW/mm²; all other traces are at approximately 13 nW/mm². WAChR variants exhibit clear and reproducible inward currents, while other tested opsins do not show a discernible response at this light level. An artifact from shutter opening immediately before photostimulation is blanked in the example traces. (B) Box plot comparing the steady-state current generated by WAChR variants and other opsins in response to irradiance of 15 nW/mm² or less. (C) A diagram of the curtain test setup for testing ambient light sensitivity, where the light-blocking curtain surrounding the patch-clamp rig is lifted to expose the cell to standard indoor office lighting. (D) Quantification of the absolute photocurrent generated by various opsins in response to ambient light exposure. The WAChR variants show the most robust responses. (E) Ambient light response from (B) normalized and displayed as a fraction of each opsin’s saturated photocurrent. Representative photocurrent traces from different opsins during exposure to ambient light (indicated by the shaded boxes). The inset traces show robust responses to direct photostimulation at an optimal wavelength (λ_max_), confirming functional expression.

Since we detected robust WAChR photocurrents at such low irradiances, we speculated that they might exhibit robust responses to indoor office light. To test this idea, we performed the simplest assay we could think of, which we term “the curtain test”: after obtaining a patch-clamp recording in dark conditions, we lifted the curtain on the patch-clamp rig, exposing the cell to ambient light from the standard indoor office lighting (still shielded from 5 sides, Fig 6C, Fig S8D). We observed robust responses to this assay from all the WAChRs that we tested (Fig 6D-F). Other frontier opsins exhibited no or comparatively weaker responses to this test (Fig 6D-F). As a positive control, we also strongly photostimulated the same cells in a subset of experiments, which confirmed that these cells were healthy and had robust opsin expression (Fig 6E and insets in Fig 6F).

To validate the activity of WAChRs in neurons, we derived human neurons from induced pluripotent stem cells (iPSCs) and transduced them with AAV-EF1a-WAChR-M219L, a variant identified earlier in the screen before WAChR-m. Consistent with the large inward currents observed in HEK cells, the curtain test resulted in a robust spiking response (2 of n=3 neurons capable of firing action potentials spiked upon curtain-opening, Fig 7A). We proceeded to inject the mouse cortex with AAV-EF1a-WAChR-m, to evaluate functionality *in vivo*.

**Figure 7.**
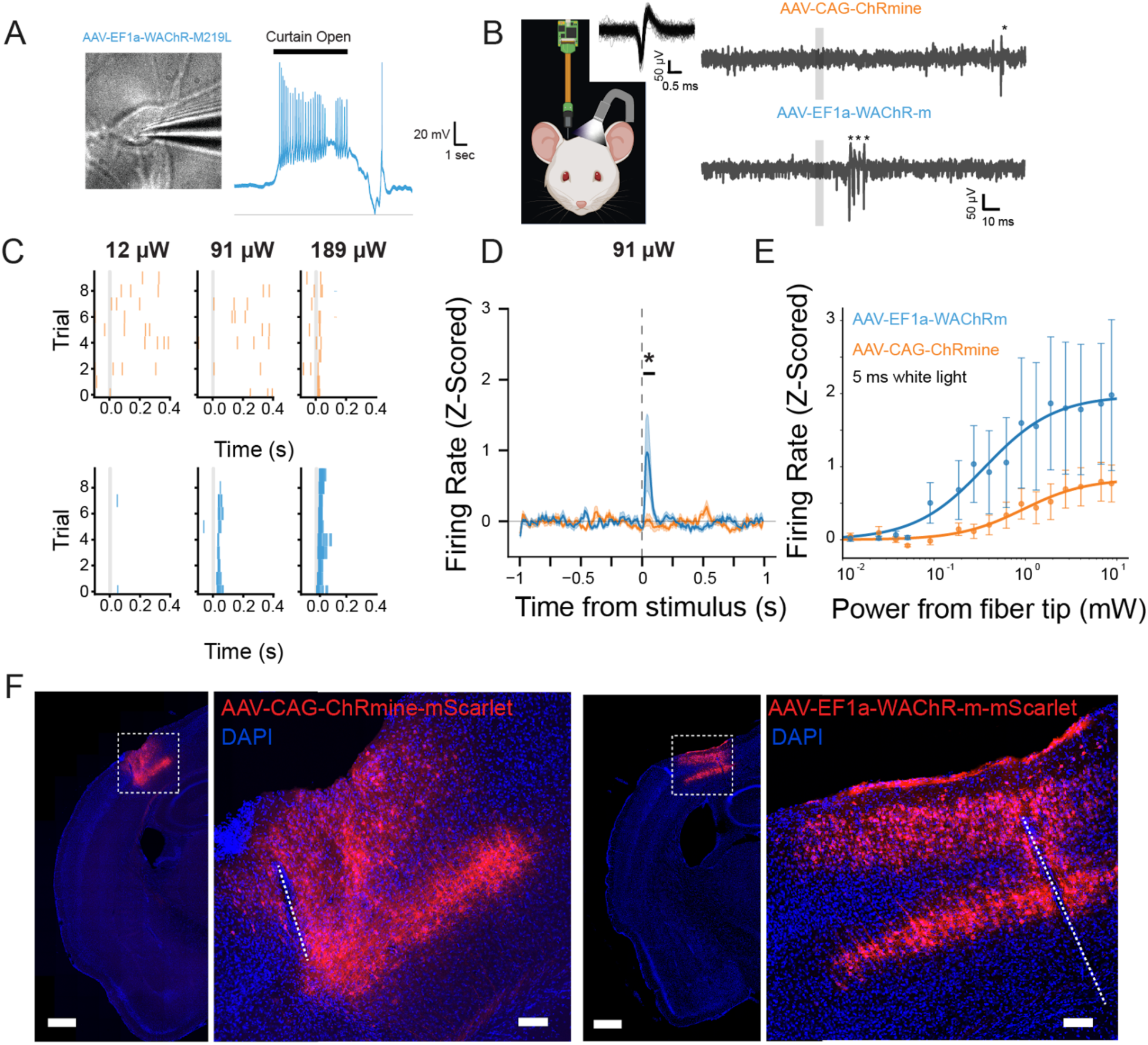
WAChRs are potent tools for optogenetic excitation (A) A human iPSC-derived neuron expressing WAChR-M219L (left) fires robustly during the curtain test (right). (B) Schematic of the *in vivo* recording setup, where a neuropixels probe records neuronal activity in an awake, head-fixed mouse expressing an opsin. Example spike waveform for one unit (inset). Example single electrode raw extracellular traces for ChRmine and WAChR-m are shown on the right during photostimulation. Spikes isolated for example clusters identified by *. (C) Raster plots showing spike times from representative units expressing either ChRmine (top, orange) or WAChR-m (bottom, blue) in response to white light pulses of increasing power. WAChR-m drives more reliable spiking across trials, especially at lower light levels. (D) The averaged, Z-scored firing rate in response to a low light power (91 µW 5ms white light pulse) is significantly greater in WAChR-m compared to ChRmine units during post light stimulation time window 2-100ms (Mann-Whitney U test, WAChR-m n=25, ChRmine n=26, U=196, * indicates p=0.015). (E) Comparison of individual firing rate (Hz) before photostim (-100 to -2ms) and after photostim (+2 to +100ms) in isolated ChRmine (orange) and WAChR-m (blue) individual units during 5 ms 91 µW photostim trials. Group means and standard error indicated in bold. WAChR-m shows greater firing rate in post stimulation window compared to ChRmine. Histological images confirm robust opsin expression in the mouse cortex for both ChRmine (left) and WAChR-m (right). Scale bars are 500 µm (large image) and 100 µm (inset). Dashed lines indicate recording probe insertion tract.

AAV-CAG-ChRmine, which is effective at optogenetically exciting neurons with low light power^7^, was injected as a positive control.

Several weeks later we recorded from the cortex proximal to the injection site with a neuropixels probe. We stimulated the brain with a white LED fiber positioned near the craniotomy (Fig 7B). We obtained stable recordings and performed optical stimulation for a variety of intensities and durations. Since the fiber was not implanted into the brain, but only placed near the probe, we reported the power of light exiting the tip of the fiber. The exact irradiance of light impinging on the neurons is not known due to variability in depth in the brain and distance from the fiber. We consistently observed significant increases in firing rates in WAChR-m mice at light conditions that did not evoke responses in ChRmine mice (Fig 7C-D).

WAChR-m was more sensitive to white light stimulation at each stimulation duration tested (5-500ms, Fig S9-10). For stimuli intense enough to elicit responses from both ChRmine and WAChR-m expressing animals, WAChR-m consistently elicited larger light-evoked responses (Fig 7E). We observed similar firing properties in mice injected with AAV-EF1a-WAChR-M219L (Fig S9-10). Histological examination of the brain identified morphologically normal neurons robustly expressing each opsin with similar laminar distribution (Fig 7F, Fig S11). This indicates that differences in activation thresholds were likely not the result of different laminar distribution of transduced neurons, and shows that long-term viral expression of WAChR-m is well-tolerated in mouse cortical neurons. Together, these data establish WAChRs as a family of potent excitatory optogenetic actuators.

## Discussion

We developed a high-throughput electrophysiology assay to measure the functional properties of channelrhodopsins. To date we have generated and characterized over 1,750 opsins, which is the largest functional opsin screen of which we are aware. For this study, we described one arm of this campaign aimed at creating highly sensitive and performant opsins. We identified the F240A mutation to WiChR, which converted it to an excitatory channel, WAChR. We further engineered this channel to alter its kinetics and sensitivity by creating and screening over 650 WAChR variants. We identified three variants, WAChR-f, WAChR-m, and WAChR-s, along the frontier of speed and sensitivity and demonstrated that they are amongst the most sensitive and efficacious optogenetic actuators described to date.

### Channelrhodopsin functional properties

Comprehensively evaluating CRs as optogenetic tools is notoriously difficult, in part because there are not universally accepted standards for characterizing an opsin. In our screen, we focused on measuring the functional properties that govern their performance under low light conditions, which bears on their potential use in vision restoration. The utility of CRs for this purpose emerges from the complex interplay between their fundamental biophysical characteristics. Their ion selectivity, in combination with the ionic concentrations in and around an expressing cell, defines a reversal potential that determines how channel opening will influence the membrane potential, while their conductance - a product of the single channel conductance and the number of CR molecules that have been produced and trafficked to the membrane - determines the magnitude of that influence. Together, these factors define the photocurrent it can generate at saturating light intensities. A CR’s sensitivity - how efficiently photons trigger channel opening - sets the minimum light requirements for activation, with the action spectrum defining which wavelengths of light are most effective at doing this. While CRs can be simplistically described as transitioning between closed and open states, their photocycles involve multiple intermediate states that govern the channel’s dynamic behavior: transitions between closed and open states create the on-kinetics and off-kinetics that determine how quickly photocurrent initiates and ceases after light onset and offset, while transitions to intermediate states are thought to underlie phenomena like desensitization and recovery that alter channel function during sustained or repeated illumination. We summarized this complexity with only a few parameters (A_sat_, K, T200_opt_, and τ_off_) which allowed us to navigate the complexity of protein function in this screen.

Measures of macroscopic photocurrent magnitude, such as A_sat_, provide useful and differentiating information about the efficacy of recorded CRs, but we expect them to vary considerably between recordings of the same CR owing to differences in the number of CR molecules being observed in each recording. Conversely, the half-saturation constant K, which is the canonical measure of sensitivity, should theoretically be independent of the number of CRs being measured in a given recording, and thus more consistent. In practice we found this to be the case, though we still observed considerable spread in our estimates of K. We also note that the maximum stimulation intensities used in this assay were approximately 1 mW/mm^2^, which is below saturation for many CRs and might limit the identifiability of these parameters, especially A_sat_. Despite these limitations, we adopted T200_opt_ as our primary measure of low-light performance in this campaign, as it integrates information about how efficiently photons can trigger CR opening with noisy but practically crucial context about how much photocurrent can be mustered by the CRs present in a cell. T200_opt_ is also readily interpretable: it indicates the amount of light an operator must deliver to a CR expressing cell in order to evoke 200 pA of photocurrent (equivalent to approximately 2.85 nS under our recording conditions). While this value does not directly translate from HEK cells to neurons, we have made the working assumption that measuring T200_opt_ in our experimental system for a given CR is a proxy for how much light that CR will require to be an effective actuator in neural applications.

### Limitations and caveats

The gold standard for CR characterization is manual whole-cell patch-clamp. While rigorous, this method is slow, laborious, and requires significant expertise, making it fundamentally unscalable for large engineering campaigns. All-optical approaches offer higher throughput but provide an indirect and at least somewhat impoverished view of a channel’s biophysical properties. Our automated patch-clamp assay was therefore designed as a pragmatic compromise, balancing the rigor of electrophysiology with the throughput required for systematic protein engineering.

Several limitations of our assays deserve further discussion. First, our photostimulation duration of 90 ms is often not long enough to reach the peak photocurrent. The time to reach this peak depends on both stimulation conditions as well as opsin properties; opsins with sufficiently slow onset kinetics are systematically disadvantaged in this context because their peak photocurrents will be clipped more frequently and/or severely by the cessation of stimulation. Second, the peak light intensity of our system is not saturating for all opsins we screened, which would lead to an underestimation of the maximal photocurrent amplitude.

Third, we stimulated quickly and repetitively (420 trials with a 3 s trial period), which means that slower dynamics like desensitization can be lost to some extent in trial-averaging. Furthermore, the interval between bouts of photostimulation is much shorter than is required for full recovery from desensitization, which disadvantages opsins that are strongly desensitizing. This experimental design was constrained by our need to obtain a sufficient number of measurements over the large number of stimulation conditions required to estimate the response surface. We also considered that these choices provide a degree of implicit selection for desirable features for vision restoration by penalizing opsins that are slow (beyond a point) or desensitizing. Our assay does not capture many aspects of CR function, including the influences of voltage, pH, and temperature on photocurrent, ionic permeability ratios, unitary channel conductance, and the hysteresis of CR photobiology on the timescale of seconds and longer. These properties are critically important to understanding CR biophysics, but comprehensively addressing the full suite of CR functional properties in a single assay is intractable due to technical complexity and the curse of dimensionality.

We opted to use plates that employ an “ensemble” recording configuration in our automated patch-clamp system, in which each recording site integrates the current from up to 20 micropipettes in parallel, meaning the measured photocurrent represents the summed activity of multiple cells. Therefore, the magnitude of the observed current is not only determined by the intrinsic properties of the opsin and its expression density within a single cell but also by the number of successfully patched cells contributing to the ensemble recording.

The number of contributing cells is likely dependent on the health of transfected cells, which can be affected by the expression of a given opsin variant. A construct that harms cell viability will result in fewer successful patches and a smaller total current, regardless of the opsin’s intrinsic properties. This effect is compounded by the population statistics of opsin expression and membrane trafficking efficiency within a given sample of transfected cells. The total photocurrent we measure is therefore a product of how many cells are recorded, how much opsin DNA is in each cell, and how well the opsin is expressed and tolerated in each one.

Finally, delivering CR DNA to HEK293T cells via lipofection introduces inherent variability. Consistent with previous reports, expression and membrane trafficking varies considerably between different CRs^5,14,17^. In some cases, we found that we were unable to achieve high quality recordings for particular constructs due to what were ostensibly failures to achieve suitable surface expression for recording. Some constructs failed to express or expressed only at low levels, while others expressed at high levels that caused aggregations and/or appeared to harm the overall health of the cells, which likely hampers the likelihood of achieving successful patch-clamp recordings. These factors contributed to a selection bias in our assay: opsins that express, fold, and traffic well in HEK293T cells via lipofection consistently perform better than those that do not. This effect is highlighted by the systematic increase of A_max_ and decrease of T200_opt_ when using AAVs relative to lipofection (Fig S6).

We proceeded with a standardized transfection protocol for several pragmatic reasons.

Developing and validating a bespoke protocol for each of the thousands of unique opsins would be intractable. Instead, we optimized a ‘one-size-fits-all’ protocol using ChRmine as a baseline. While this approach likely produced false negatives, meaning that we might miss or underestimate the performance of some opsins, it was essential for enabling a large-scale campaign. The goal was not to perfectly characterize every opsin, but rather to efficiently identify promising candidates for further engineering.

### Mechanism of extraordinary sensitivity

Enhancing the low light sensitivity of CRs is a long-time goal of the field. Some earlier studies succeeded by decelerating photocurrent decay kinetics. In particular, mutations to the “DC gate” and other residues of *Chlamydomonas reinhardtii* channelrhodopsin 2 and other green algal CRs can slow channel closing by several orders of magnitude, resulting in highly sensitive “step-function” optogenetic tools which can be toggled into a long-lasting open state with a brief pulse of light^20,21,35,49,69^. Analogous mutations render the PLCR ChRmine nonfunctional^40^ and are tolerated by HcKCR1^70,71^ but might result in leak currents independent of photostimulation ^72,73^. The slow kinetics of resulting tools constrain how they can be deployed, though some studies have identified mutations that result in similar but less dramatic effects^23,32^.

Another point of attack has been the desensitization dynamics of CRs. Desensitization varies to a large degree among naturally occurring CRs, and many of the natural CRs that have been repurposed for optogenetics recently undergo much less desensitization than the first generation of optogenetic tools^8,9,11,74–77^. The mechanisms of desensitization are incompletely understood and may differ fundamentally among CRs^13,78,79^, but involve the transition of CRs into long-lived, non-conducting or weakly conducting states during sustained illumination.

Some efforts to improve CRs for low-light applications have found success in diminishing the extent of desensitization^1,32,45,80^. This might reflect changes in the kinetics of photocycle transitions that favor the accumulation of the primary open state, similar to step-function opsins, but in principle could also be accomplished by increasing the conductance of secondary open states.

The remarkable sensitivity of the WAChR variants likely stems from structural and photocycle dynamics inherited from its PLCR parent opsin, WiChR. The mechanisms for high sensitivity in the PLCR family are an active area of investigation. For instance, structural studies of the related potassium channel HcKCR1 suggest that its channel opening involves minimal, energy-efficient sidechain movements rather than large-scale backbone rearrangements, which could contribute to a lower activation energy and thus higher sensitivity to light^70^. A similar mechanism appears to contribute to the high sensitivity of the PLCR GtCCR4, perhaps enabled by a unique bend in its 6th TM domain^13,75^. It is plausible that these or similar mechanisms contribute to the high sensitivity of WiChR and WAChR. We note that, under our experimental conditions, WAChR and most of its variants undergo only minimal desensitization (though in some variants the degree of desensitization was substantially greater; data not shown). However, WiChR and HcKCR1 are known to undergo a change in ion selectivity during sustained photostimulation^81^ in which the permeability of Na^+^ increases relative to K^+^, suggesting the presence of at least two open conformations in these KCRs. One possibility is that the secondary open state(s) of WAChR might remain highly conductive, but this can only remain speculative pending detailed structural and biophysical investigation.

### Applications of WAChR variants

Numerous CRs have demonstrated efficacy for vision restoration in animal models of inherited retinal disease. We were able to obtain robust recordings from several CRs promulgated for vision restoration (ChReef, ex3mV1Co, CoChR-3M, ChRmine, and ChroME2s)^33,34,45,67^. WAChR variants appear to be among the most sensitive excitatory CRs to date, and we expect that they will also perform well in preclinical models of vision restoration. Many attempts at optogenetic gene therapies have required the use of assistive devices^4,82,83^ that provide enhanced illumination to the eyes owing to the lack of sensitivity of the opsin to indoor lighting. Phototoxicity concerns from this illumination have driven a general preference for red-shifted opsins in vision restoration. However, if the frontier WAChR variants are able to drive light perception under ambient lighting conditions or with consumer virtual/augmented reality displays, then phototoxicity concerns are no longer paramount, obviating the need for red-shifting. For experimental applications requiring deep tissue penetration or multiplexing with blue-light activated sensors, other opsins are likely more suitable than WAChR variants.

While WAChR variants robustly respond to ambient office light via the “curtain-test”, a basic sanity check that we have not seen previously reported, we should bear in mind that the light levels incident on the retina are dimmer than the room due to the numerical aperture of the pupil. Once sufficient sensitivity has been achieved, speed is the other major consideration for engineering CRs for vision restoration. It is not obvious *a priori* how fast a CR must be for this purpose, but all other things being equal, faster is likely better. Notably, even WAChR-s exhibit substantially faster decay kinetics than ex3mV1Co and CoChR-3M, while exhibiting larger responses to dim light. Future experimentation with WAChR-f, WAChR-m, and WAChR-s will allow the discovery of the optimal speed-sensitivity tradeoff for vision applications. This family of ambient light sensitive opsins should be broadly useful for neuroscience applications.

## Methods

### Opsin Scaffolds, Molecular Biology, and Plasmid Construction

Opsin Scaffolds and Starting Sequences: A variety of opsin sequences served as scaffolds for this engineering campaign. The wild-type DNA sequence for *Wobblia lunata* inhibitory channelrhodopsin (WiChR) ^50^ was sourced from NCBI accession PRJDB4369_DN2329_c0_g1_i10.

Expression Plasmids (for HEK293T cell expression): Plasmid backbones containing either an EF1a or a TRE (Tetracycline Response Element) promoter were used to drive expression of all opsin variants in HEK293T cells. All plasmids conferred resistance to carbenicillin. Plasmid backbones were obtained from VectorBuilder.

Fusion Protein Constructs (for HEK293T cell expression): To facilitate membrane trafficking, localization, and visualization, the expressed opsin sequences were part of a fusion protein. The constructs, in N-terminal to C-terminal order, typically concatenate the following elements: the opsin coding sequence, a short linker (e.g., AAA), a Kir2.1 Golgi trafficking sequence (KSRITSEGEYIPLDQIDINV), the mScarlet fluorescent protein (VSKGEAVIKEFMRFKVHMEGSMNGHEFEIEGEGEGRPYEGTQTAKLKVTKGGPLPFSWDILSPQFMYGSRAFTKHPADIPDYYKQSFPEGFKWDRVMNFEDGGAVTVTQDTSLEDGTLIYKVKLRGTNFP PDGPVMQKKTMGWEASTERLYPEDGVLKGDIKMALRLKDGGRYLADFKTTYKAKKPVQMPGAYNVDRKLDITSHNEDYTVVEQYERSEGRHSTGGMDELYK) for identifying expressing cells, a Kv2.1 soma-targeting sequence (QSQPILNTKEMAPQSKPPEELEMSSMPSPVAPLPARTEGVIDMRSMSSIDSFISCATDFPEATRF), and a Kir2.1 endoplasmic reticulum (ER) export sequence (FCYENEV) (Supplementary Fig 1H) ^17,46,47,55,84^. In some cases, we also prepended the LucyRho tag to the N-terminus of opsins to attempt to improve expression and/or membrane trafficking ^58^.

### Variant Generation for Autopatch Screening

DNA Synthesis: DNA encoding designed variants was clonally synthesized commercially (Twist Biosciences, South San Francisco, CA). Generally we adopted the coding sequence of opsins that had previously been validated, however in cases where novel channelrhodopsins were being tested then human codon optimization was performed using DNAChisel^85^ or BaseBuddy^86^.

Mutagenesis: One round of random mutagenesis via error-prone PCR and one round of site-directed mutagenesis was performed on WAChR variants. The random mutagenesis round utilized the primers WiChR_F240A(pVBRP-EF1a)_F and WiChR_F240A(pVBRP-EF1a)_R (Supplementary Table 1) to amplify the WAChR coding sequence (Supplementary Table 2) using an error-prone polymerase (Agilent GeneMorph Polymerase), prior to being Gibson assembled into plasmids (NEB HiFi DNA Assembly Kit). The site-directed mutagenesis round was performed on a pool of 14 WAChR variants (WiChR_T127V_F240A, WiChR_L154T_F240A, WiChR_A158G_F240A, WiChR_N117L_F240A, WiChR_D47C_F240A, WiChR_W120Q_F240A, WiChR_W120A_F240A, WiChR_S161A_F240A, WiChR_M219L_F240A, WiChR_T127S_F240A, WiChR_G56L_F240A, WiChR_T95A_F240A, WiChR_T127V_C201A_F240A, WiChR_H229E_F240A) targeting 8 total sites: A240, F239, W120, M219, T127, C250, T252, I131. Site directed mutagenesis primers were designed using the “small intelligent trick”^87^ and are shown in Supplementary Table 1. Site-directed mutagenesis PCRs were performed with a high fidelity polymerase (Takara Bio GXL Polymerase) in one fragment before being circularized into plasmids (NEB KLD Enzyme Mix). In both mutagenesis rounds, assembled plasmids were transformed into an *E. coli* cloning strain (NEB 5-alpha) and plated onto LB agar plates containing 100 µg/mL carbenicillin. Plates were incubated overnight at 37°C and colonies were picked into 80 µL of LB with 100 µg/mL carbenicillin into 4x 384-well plates by an automated colony picker (QPix 420, Molecular Devices). Seed cultures were incubated overnight at 37°C, after which a glycerol stock (40 µL 60% glycerol + 40 µL culture) and boil prep (20 µL dH2O + 20 µL culture, then incubated at 95°C for 10 minutes) was prepared from each overnight culture. A 10X miniaturized library prep was performed on each boil prep with the Enzymatic Fragmentation Kit 2.0 and HT Universal Adapters (Twist Biosciences) before being pooled and sequenced on a NextSeq 1000 (Illumina). Basecalling and demultiplexing were performed with BCLConvert v4.2.7. Plasmid alignment and variant analysis were performed by nf-core Sarek v3.4.2^88–90^. Plasmids that contained valid and unique coding sequences were re-inoculated from the glycerol stock to be miniprepped on a Kingfisher Flex (Thermofisher) using magnetic beads (Magjet Plasmid DNA Kit, Thermofisher).

### Computational and Machine Learning-Guided Design

Overview: Our approach to designing and selecting mutants to test evolved over the course of this campaign, which was one arm of a larger opsin engineering campaign. The full details of this larger campaign are beyond the scope of this study, but we describe our methodology in brief here. Generally, our design approach went through three main stages: generation, filtering, and selection. A variety of methods were applied for generating candidates including: structure-guided hypotheses, structure-guided recombination, zero-shot mutation effect prediction with protein language models, ancestral sequence reconstruction, gradient-based markov chain monte carlo (MCMC) sampling, *de novo* sequence design, and genome mining. Our filtering pipeline proceeded through two stages: fast and slow filters. Fast filters were generally sequence-based metrics that served as a preliminary large scale filter. Candidates that passed the fast filters had their structures predicted and subsequently passed through slow structure-based metrics. Lastly, we ranked candidates based on their sequence-to-function model score and uncertainty. Candidates were synthesized and tested in batches where the subsequent results were used to re-train our sequence-to-function model.

### Generation

Genome Mining: A database of opsin sequences was generated by a protocol adapted from Zheng et al.^91^ Briefly, a seed file containing representative channelrhodopsin sequences was used to query Uniclust30^92^ with HHBlits^93^. An HMM profile is produced from the hits and used to search the following genomic/metagenomic databases with HMMER3^94^: MGV^95^, GPD^96^, SMAG^97^, JGI-UMAG^98^, IMG/VR^99^, MERC^100^, Uniref100^101^, SRC^100^, Metaclust^102^, Mgnify^103^, and JGI-IUIG^98^. Sequence hits were collected and deduplicated with MMSeqs2^102^. We additionally tested a few genome mined proteins by filtering for diversity using hhfilter^93^.

Fine-tuning PLMs: We used low-rank adaptation (LoRA)^104^ to fine-tune pretrained protein language models (PLMs) on a masked-language modeling task, with our genome mined opsin database as the training data. As base models, we used the 650M parameter version of ESM^105^ as well as its structure-aware derivative ISM^106^ and the convolutional model CARP^107^.

Zero-shot prediction of mutation effects: We used the base and fine-tuned PLMs to evaluate the approximate pseudo log likelihood of sequences near the wildtype (typically Levenshtein distance of 3 or less), using the efficient implementation described in^108^. We then calculated the log odds ratio of variant sequences versus the wildtype, since these scores amount to zero-shot predictions about whether edits will be beneficial or deleterious^109^.

One way we used these scores was to perform a “zero-shot substitution scan” across WAChR to propose candidate sequences to test. For each position in the protein, we computed log odds ratios for all 19 possible amino acid substitutions, generating a position-by-substitution matrix of predicted mutational effects. This allowed us to visualize the distribution of single-step edits from any given sequence, revealing both highly constrained positions (where all substitutions showed negative log odds) and potentially mutable sites (where one or more substitutions showed positive log odds). We selected candidate sequences from these using a breadth-focused strategy, sampling mutations across many positions throughout the protein rather than exhaustively exploring individual sites. For positions which had substitutions with favorable log odds ratios, we selected one of the top-scoring mutations (or occasionally the 2-3 best mutations when multiple substitutions showed similarly high scores).

Ancestral Sequence Reconstruction: We leveraged ASR in order to try and generate variants with improved stability. Briefly, a multiple sequence alignment (MSA) of well-characterized channelrhodopsins was generated with MAFFT^110^. The MSA was manually curated before being fed to the ancseq pipeline^111^ which generates a maximum likelihood phylogenetic tree with IQTree2^112^. Ancseq then uses the phylogenetic relationships to infer the marginal posterior probabilities for each residue at each ancestral sequence node. In addition to taking the maximum *a posteriori* (MAP) sequence from a chosen ancestral node, additional candidate ancestral sequences are sampled from its posterior distribution (to take into account the uncertainty within the phylogenetic inference) using the minimum posterior expected error (MPEE) criteria^113^ as follows:

1. If a residue has a gap in the MAP, then all probabilities for residues at that site are set to 0. Otherwise, the probability of a gap at that site is set to 0.
2. The MPEE set is calculated for each site.
3. Then the probability for each residue that is not in the MPEE set at each site is set to 0.
4. The remaining residue probabilities are re-normalized to sum to 1.
5. A large number of samples are drawn from the MPEE-normalized posterior distribution of the ancestral sequence.
6. The top k sequences with highest posterior probability are selected as candidates.

This sampling procedure additionally took inspiration from Eick et al. 2012 and 2017^114,115^. The idea is to avoid pathological residues while sampling the phylogenetic distribution.

Recombination: Many previous opsin engineering campaigns have utilized recombination of existing opsins to generate new and diverse functionality. We included additional variants in our screen that were composed of recombined opsins using junctions that were adapted from Bedbrook et al.^15,116^ The swappable block junctions were based on the 10 contiguous block strategy and modified such that there were 6 swappable blocks that did not change the native multimeric interface of an acceptor opsin (which we chose to be WAChR-M219L), and furthermore were shifted if possible such that the junction occurred within one of the 7 transmembrane helices. This was done with the hypothesis that recombination events within conserved secondary structure elements are less likely to disrupt protein activity.^117^ Donor opsins included HcKCR1, HcKCR2, GtCCR4, ChRmine, HulaCCR, B1ChR2, and WlChR2-4. A combinatorial library of chimeric opsins from these parts was generated.

*De novo* Sequence Design: We generated an additional set of candidates using ProteinMPNN^118,119^. We sampled sequences conditioned on predicted structures of WAChR as well as other chimeric opsins.

Gradient-Based MCMC Sampling: To conditionally sample from our sequence-function model, we utilized the EvoProtGrad framework^120^ to perform Gradient-Based MCMC Sampling. This method combines multiple distinct signals: the gradient from our sequence-to-function model(s) to guide sampling toward improved activity, and the gradient from the base ESM2 model to ensure that generated sequences maintain high evolutionary likelihood.

Rational design: We used information from structural predictions, MSAs, PLM likelihood scans, and existing literature to select mutations and regions of interest.

### Filtering

Sequence-to-Function Model: We developed a machine learning framework to predict opsin properties from sequence and experimental context. Our training dataset combined our internally-generated measurements with data ingested from public sources, providing diverse coverage of sequence-to-function relationships across different experimental conditions. To evaluate model generalization, we employed multiple strategies for generating training/validation folds: (1) random splits grouped by protein identity; (2) sequence-similarity based splits using MMseqs2 clustering to group related variants; (3) temporal splits based on when each opsin first appeared in our experimental pipeline, allowing assessment of prospective prediction performance. We trained ensemble models across these different fold strategies to obtain robust predictions.

We experimented with a number of different modeling strategies over the course of the campaign, which we describe here in brief. In the early phases of the campaign, we primarily used LoRA fine-tuning to train single-task regression, classification, or Bradley-Terry regression^121^ models, typically using targets that were aggregated at the level of opsins or constructs.

Later, we adopted a multi-task architecture. The multi-task model operated on individual assay measurements rather than aggregated protein/plasmid-level statistics. For each observation, we embedded the opsin sequence using either base or fine-tuned PLMs. A text representation of experimental context was encoded separately using Bio-TinyBERT^122^, which included information such as cell line, expression protocol parameters, promoter, and accessory protein elements such as trafficking signals or fusion partners. These dual embeddings were integrated through a cross-attention mechanism, allowing the model to condition predictions on experimental setup.

The training objective combined multiple complementary losses to leverage different views of the data. For regression tasks (λ_max_, τ_off_, A_max_, K, and T200_opt_), we used a Bradley-Terry pairwise ranking loss^121^ learned from relative property comparisons between opsin pairs, weighted by sequence similarity to reward correct ranking of closely-related variants while preventing mode collapse; Huber regression loss provided direct property prediction robust to outliers; auxiliary classification losses on discretized property bins at multiple resolutions captured ordinal relationships; and contrastive loss with momentum contrast^123^ learned property-preserving representations. For binary classification tasks (functional vs nonfunctional, expressing vs non-expressing) we used cross-entropy classification losses as well as contrastive loss.

To balance competing objectives, we employed gradient-based multi-task balancing using GradNorm^124^ to dynamically adjust task weights based on relative training rates, combined with PCGrad^125^ to project conflicting gradients and prevent negative transfer between tasks. During training, we also incorporated augmentation strategies including sequence masking and dropout of some or all of the contextual embeddings.

Candidate Filtering Pipeline: Every candidate opsin that we generated proceeded through a fast and then slow filtering stage. The fast filtering stage utilized several common protein design metrics^126^ including: PLM log-likelihood, out-of-distribution filtering^127^, distance metrics to a referent protein (typically WAChR) for PLM embeddings as well as membrane topology predictions via TMbed^128^ and hard-coded functional constraints (e.g. K251 in WiChR was constrained because it is the Schiff base lysine). The fast filtering stage additionally used inference outputs from our sequence-to-function model ensembles described above.

The slow filtering stage used a local version of ColabFold^129–131^ (later Boltz2^132^) to predict the protein structure of candidates that passed the fast filtering stage. The predicted structure was used to calculate metrics of pLDDT and pTM. We passed these structures back into ProteinMPNN and/or LigandMPNN in order to calculate sequence recovery and self-consistency perplexity^133^.

### Selection

Following the hard-filtering stage, we used a subset of the same metrics for ranking and selection of variants. To guide sequence selection, we used an acquisition function based on either the Upper Confidence Bound^134^ or Noisy Expected Hypervolume Improvement^135^. In some cases we incorporated additional terms to promote diversity within a particular batch by greedily penalizing selection of new variants for being close in embedding space to previously selected variants.

As with other parts of the methodology, our selection strategy was fluid throughout the campaign. Within a given round of synthesis, we often generated several batches of designs using different selection criteria. Early on, we prioritized broad exploration and relied heavily on PLM-likelihood based guidance, but shifted towards using inference from our sequence-to-function ensembles later as more training data became available. Our objectives for the campaign also evolved as we gained a more mature understanding of our capabilities. For instance, we noticed relatively early that we achieved better performance at predicting τ_off_ than other metrics (data not shown). This likely reflects the fact that τ_off_ measurements appeared to be more precise than our other measurements (Fig. 4B). To leverage this, we increased our emphasis on improving kinetics while also improving or at least preserving sensitivity and devoted more of our sample budget to reflect this. We also made affordances to the human-in-the-loop nature of designing and running protein engineering campaigns; namely, we allowed ourselves an ‘override’ option to include particular designs (e.g. for a rational design hypothesis) in synthesis batches, which was purely a concession to our own curiosity.

### Cell Culture and Transfection (for HEK293T cell experiments)

Cell Line and Culture Conditions: Human Embryonic Kidney 293T (HEK293T; ATCC, Manassas, VA) cells were used for the heterologous expression and electrophysiological screening of all opsin constructs. Cells were maintained in Dulbecco’s Modified Eagle Medium (DMEM) supplemented with 10% Fetal Bovine Serum (FBS), 1% GlutaMAX, and 1% Penicillin-Streptomycin (referred to as DMEM++) in a humidified incubator at 37°C and 5% CO2. Tissue culture plates (6-well or 96-well) were coated with 1X GelTrex LDEV-Free hESC-Qualified Reduced Growth Factor Basement Membrane Matrix (Thermo Fisher Scientific) prior to cell seeding. For experiments, 6-well plates were seeded with 1.2e6 cells per well, and 96-well plates with 3e4 cells per well.

Transient Transfection Protocol: HEK293T cells were transiently transfected using Lipofectamine 3000 Kit (Thermo Fisher Scientific) in OptiMEM I Reduced Serum Medium (Thermo Fisher Scientific), according to the manufacturer’s instructions. For 6-well plates, cells were typically transfected with 500-1000 ng of plasmid DNA for EF1a promoter constructs or 2500 ng for TRE promoter constructs. Cells were used for patch-clamp recordings 24-48 hours post-transfection. For constructs under a TRE promoter, expression was induced by adding doxycycline (Sigma-Aldrich, St. Louis, MO) to the culture medium at a final dilution of 1:6500 from a stock solution (20mg/mL), approximately 24 hours after transfection.

Automated Patch-Clamp Electrophysiology

An automated, high-throughput patch-clamp assay was performed using a modified IonFlux Mercury 16 system (Cell Microsystems). To accommodate the custom light path, the instrument’s standard heater was removed, and the IonFlux unit was elevated on metal posts: For the standard autopatch assay, photostimulation was delivered from underneath the IonFlux plate. Light from a Lumencor SPECTRA X light engine (Lumencor Inc.) was transmitted via a liquid light guide. The light path was further modified with a condenser lens to expand the beam. A galvo-galvo scanning system (Thorlabs) was used to scan the light between the 8 recording sites on the IonFlux plate. Light stimuli were typically 90 ms in duration, with trials run on a 3-second period. The standard assay typically consisted of 7 wavelengths (440, 475, 510, 555, 575, 637, and 748 nm), each tested at 10 different irradiances, with 6 repetitions per condition, totaling 420 trials per recording site. Light intensities were varied, ranging from approximately 1 μW/mm² to 1 mW/mm². For some channels, in order to achieve low irradiances, pulse width modulation (1 ms period, duty cycles ranging from 5-95%) was implemented by gating the output of the Lumencor using a 200kHz digital line. Irradiances were calibrated at the sample plane using a Thorlabs photodiode (e.g., S120VC or S130VC) with a 1 mm diameter pinhole attached. Control of the Lumencor light engine and the galvo system was achieved using a National Instruments DAQ (NI-DAQ) board, which received trigger signals from one of the compound wash wells of the IonFlux system to synchronize light delivery with the electrophysiology protocol using custom software. Cells were voltage-clamped at a holding potential of -80 mV. Data was collected at 2kHz.

Cell Preparation: Prior to autopatching, HEK293T cells were washed twice with 1X Dulbecco’s Phosphate-Buffered Saline (DPBS), then detached from the culture plate using Accutase (Innovative Cell Technologies) by incubating at 37°C for 10 minutes. Cells were collected, pelleted by centrifugation at 200xg for 1 minute, and resuspended in external solution to a final concentration of approximately 8.0 x 10^6^ cells/mL (or a minimum volume of 250 µL if cell yield was lower). The system utilized IonFlux’s SBS-standard 96-well ensemble plates. Appropriate wells were filled with external and internal solutions.The external solution consisted of (in mM):150 NaCl, 4 KCl, 2 CaCl2, 2 MgCl2, 10 HEPES, 10 Glucose; pH adjusted to 7.4 with NaOH, 310mOsm. The internal solution (in mM): 140 K-Gluconate, 10 EGTA, 2 MgCl2, 10 HEPES pH adjusted to 7.2 with KOH, 290mOsm.

Plate Reuse Protocol: To maximize throughput and minimize consumable costs, a robust plate cleaning and reuse protocol was implemented. Following an experiment, any remaining liquid was discarded. The washing procedure involved sequential treatments: 250 µL of deionized H_2_O was added to all wells (except ‘OUT’ wells of each section). The plate was centrifuged at 500xg for 5 minutes. The liquid was discarded, and this water wash and spin step was repeated once. The deionized water wash and spin steps were repeated using a 0.01 g/mL (1% w/v) solution of Tergazyme™ Enzyme-Active Powdered Detergent (Alconox, White Plains, NY). The deionized water wash and spin steps were repeated using a 10% bleach solution. Plates were rinsed three times with deionized H_2_O using a CombiDrop multichannel dispenser (400 µL/well). Following this, deionized H_2_O was added to all wells (except “OUT” wells), and the plate was centrifuged at 500g for 5 minutes; this step was repeated once. If a plate was to be used later the same day, an additional water washing step was performed using the IonFlux system, and the plate was then stored at 4°C. If not being used the same day, the plate was stored at 4°C after the final CombiDrop/centrifugation water rinses, and a water washing step using the autopatch system was performed no more than 24 hours before its next use.

### Manual Whole-Cell Patch-Clamp Electrophysiology

#### HEK293T recordings

For whole-cell patch-clamp recordings, opsin-expressing HEK293T were seeded onto poly-d-lysine coated glass coverslips (Neuvitro). Patch pipettes (4-6MOhm) were pulled from borosilicate glass (G150TF-4; Warner Instruments) and filled with internal solution containing (in mM): 123 K-gluconate, 10 EGTA, 4 NaCl, 2 MgCl2, 10 HEPES, pH adjusted to 7.2 with KOH, 290mOsm. Coverslips were placed in external solution containing (in mM): 120 NaCl, 4 KCl, 2 CaCl2, 2 MgCl2, 10 HEPES, 40 Glucose, pH adjusted to 7.2 with NaOH, 320mOsm. Reported membrane potentials are LJP-corrected.

Cells were held at -70mV in voltage clamp recordings. No current was injected in passive current clamp recordings. Recordings were using a Digidata 1550B digitizer and Multiclamp 700B (Molecular Devices) amplifier and sampled at 10kHz. All data was acquired using pCLAMP software (Molecular Devices, version 11.4).

Precise light dose response curves and action spectra were generated by stimulating cells with varying irradiances and wavelengths of light in dark-adapted conditions. For action spectra, photocurrents (initial slope, peak, and steady-state components) were measured in response to light stimuli of varying wavelengths delivered by TLS130B-300X tunable light source (Newport Corporation, Irvine, CA) coupled to a microscope (SOM, Sutter Instruments) . Photocurrents were subsequently normalized by the measured photon density for each specific stimulus condition to determine the relative efficacy of different wavelengths. Opsin expressing cells were visualized using IR camera (Thorlabs) and IR condenser light source (Olympus). Irradiance was modulated using a motorized filter wheel containing neutral density (ND) filters. Different sized slits installed in the monochromator (e.g., 280 µm, 1240 µm, 5 mm) were used to balance maximum power output and spectral resolution depending on the experimental requirements. The large majority of experiments were performed using the 5mm slits (“broad-band configuration”) which generates light with a spectral full-width half maximum (FWHM) of approximately 47nm (measured at target wavelength of 510 nm) with most of the remaining experiments performed in the 280 µm (“narrow-band configuration”) which had an FWHM of approximately 7nm. Light power was measured using Thorlabs photodiode and spectral output using Ocean Optics spectrometer. Irradiances for a given stimulation condition depended on both the attenuation (set by the ND filter) and the wavelength (because of the spectrally non-uniform output of the arc lamp). For experiments shown in Fig 5A (broad-band configuration), irradiances for 480 nm stimulation (WAChRs and CoChR-3M) ranged from ∼270 nW/mm^2^ to 2.6 mW/mm^2^; 510 nm (ChroME2s): ∼190 nW/mm^2^ to 2.4 mW/mm^2^; 525 nm (ChRmine and ChReef): ∼190 nW/mm^2^ to 2.2 mW/mm^2^; 540 nm (ex3mV1Co): ∼170 nW/mm^2^ to 2.1 mW/mm^2^. See also Fig S7A,B.

Photostim was 500ms-1s with 10s ITI for dose response assays and 300ms with 1.2s ITI for action spectra. For ambient light stimulus recordings (the “curtain test”), the light barrier was briefly lifted, held open for ∼1-2 seconds, and then returned to a closed position. Irradiances measured at the sample were typically in the range of 500 nW/mm^2^ to 1μW/mm^2^ when the light barrier was lifted and ∼2-5 nW/mm^2^ when the barrier was closed.

#### iPSC-Derived Neuron Recordings

Differentiated iPSC-derived neurons ^136^ were seeded onto poly-d-lysine coated coverslips and transduced with 10k MOI of AAV2.7m8 EF1a>WAChR(M219L)-G-mScr-SE. Neurons were 3-4 weeks old at the time of whole-cell patch-clamp recording. Recordings were acquired as above with external solution containing (in mM): 150 NaCl, 4 KCl, 2 CaCl2, 2 MgCl2, 10 HEPES, 10 Glucose; pH adjusted to 7.4 with NaOH. Internal solution contained (in mM): 135 K-gluconate, 8 NaCl, 4 MgATP, 0.3 Na3GTP, 0.3 EGTA, 10 HEPES, pH adjusted to 7.2 with KOH. Current was injected to hold neurons at -70mV if different from resting membrane potential. For ambient light stimulus recordings (the “curtain test”), the light barrier was briefly lifted and replaced.

#### *In Vivo* Extracellular Electrophysiology

All animal procedures were conducted in compliance with Science Corporation’s Animal Care and Use Committee. Adult (typically 8-12 weeks old) male and female C57BL/6J mice (Charles River) were used for *in vivo* experiments. Mice were group-housed with ad libitum access to food and water on a standard 12:12 light:dark cycle. Positive control groups (n = ∼4 mice) were injected with AAV-CAG-ChRmine. Experimental groups (n = ∼8-12 mice) were injected with AAV-WAChR variant(s).

#### Stereotaxic Virus Injection

Mice were anesthetized with isoflurane (2%), and administered 2 mg/kg of dexamethasone as an anti-inflammatory and 0.05 mg/kg Ethiqa as an analgesic (2% in oxygen) and placed in a stereotaxic frame (Kopf Instruments). A small craniotomy (e.g., ∼0.5-1 mm diameter) was made over the target injection site, parietal association cortex outside of primary sensory cortex (LPtA1, coordinates relative to bregma: -2.0 mm Anterior/Posterior, +/-2.0 mm Medial/Lateral, -0.3 mm Dorsal/Ventral from pia). AAVs were injected using a borosilicate glass micropipette (tip diameter ∼10-20 µm) connected to a microinfusion pump (Nanoject III, WPI). Typically, 100-300 nL of virus (target titer E11 vg/mL) was injected per site at a rate of 50 nL/min. The pipette was left in place for 5-10 minutes post-injection before slow withdrawal. Skin was sutured, and animals received post-operative analgesia. Mice were allowed to recover for 2-4 weeks for optimal opsin expression.

#### AAV Viral Constructs

The following adeno-associated virus (AAV) vectors were used:EF1a>{WAChR-M219L}-GFSE:WPRE3; EF1a>{WAChR-s}-GFSE:WPRE3; EF1a>{WAChR-m}-GFSE:WPRE3; EF1a>{WAChR-f}-GFSE:WPRE3; EF1a>{ChReef}-GFSE:WPRE3; EF1a>{ex3mV1Co}-F:WPRE3; EF1a>{CoChR-3m}-F:WPRE3; EF1a>{WAChR-m}-GSE-HA-P2A-GFE:WPRE3; CAG>ChRmine-GFSE:WPRE3; CAG>ChroME2s-GFSE:WPRE3.

G refers to the Kir2.1 Golgi trafficking sequence, F refers to mScarlet, S refers to the soma-targeting sequence from Kv2.1, and E refers to the Kir2.1 endoplasmic reticulum export sequence. We omitted our standard signal peptides from EF1a>{ex3mV1Co}-F:WPRE3 and EF1a>{CoChR-3m}-F:WPRE3 in order to more closely reproduce the reagents from previous reports on these opsins^33,34,137^ where the opsins were fused only to a fluorophore; however, we replaced the fluorophores used in these reports (Venus and GFP) with mScarlet. While most constructs used the EF1a promoter, we chose to use pre-existing vectors incorporating the CAG promoter for ChRmine and ChroME2s as a matter of cost and expedience; we recognize that this is less than ideal.

For in vivo cortical virus injection, viruses were diluted in sterile saline to a target titer of 1e11 vg/mL. For HEK293T cell transduction, 10-20e4 MOI of virus was added to cell media 1-4 days prior to patch-clamp recording. All constructs were packaged commercially (VectorBuilder Inc.) in the AAV2.7m8 serotype.

Cranial Window and Headbar Implantation

Approximately 1-2 days prior to *in vivo* recording, mice underwent a second surgery. Under anesthesia, a ∼1 mm x 1 mm cranial window was created over the injection site. A custom-designed headbar was affixed to the skull using resin (RelyX Unicem, 3M). A perimeter of resin was built around the craniotomy to create a well for artificial cerebrospinal fluid (ACSF) during recordings. The cranial window was covered with an opaque silicone elastomer (Kwik-Sil, WPI).

#### In vivo Electrophysiology Recordings

Awake, head-fixed mice were positioned on a running wheel in an optically isolated chamber. Kwik-Sil was removed and the cement well was filled with ACSF. A silver pellet electrode was placed in the ACSF and the recording probe (Neuropixel 1.0) was slowly lowered to final recording depth of 1500um below cortical surface at a rate of 100um every 1-2 minutes. Probe placement was allowed to stabilize for 10-15 minutes prior to onset of recording. Data was sampled at 30kHz and light was delivered via an optical fiber (e.g., 200 µm core diameter, NA 0.22-0.39) coupled to a white light LED (Thorlabs), positioned at the cortical surface. Light power was measured at the fiber tip using a photodiode power meter (Thorlabs PM100D with S130C sensor). For each power setting, white light stimulus pulses (5 or 500 ms pulse width) were delivered followed by a 5-second fixed inter-trial interval. TTL pulses from the stimulus generator (MCC DAQ) synchronized light delivery with electrophysiological data acquisition in SpikeGLX.

### Data Analysis

Electrophysiological data were analyzed using custom and commercial software. To characterize opsin responses from automated patch-clamp data, peak photocurrent amplitudes (y) were modeled as a function of photostimulation irradiance (E) and wavelength (λ) using a two-stage model. The model was fitted to the median of the maximum peak photocurrent amplitudes recorded for each wavelength and irradiance condition (6 repetitions per condition).

Stage 1: Action Spectrum Modulation: The effective irradiance (E_λ_) at a given wavelength was modulated from the delivered irradiance (E) based on a Gaussian approximation of the opsin’s action spectrum:

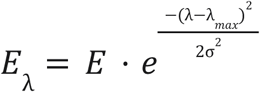

where λ_max_ is the wavelength of maximum opsin sensitivity and σ is the standard deviation defining the breadth of the action spectrum.

Stage 2: Hill Response Function: The modulated irradiance (E_λ_) was then used as input to a standard Hill equation to model the photocurrent response:

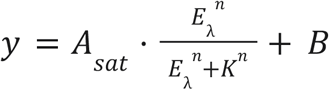

where A_sat_ is the maximum photocurrent amplitude, K is the half-saturation constant (irradiance required to reach half-maximal response, sometimes referred to as effective dose 50 aka ED50), n is the Hill coefficient (slope of the dose-response curve), and B is the baseline current. For each opsin variant screened via automated patch-clamp, the parameters λ_max_, σ, A_sat_, K, n, and B were fitted to the experimental data using a 5-fold cross-validation procedure. Fit quality was evaluated using the root mean squared error and R² for hold out data in each fold. The final reported parameters for an opsin variant represent the average of the parameters obtained from these splits.

Prior to fitting and inclusion in the dataset, all recordings underwent quality control. This involved visual inspection of each recording, with the option to flag and exclude assays exhibiting anomalies. Most commonly, this took the form of a reduction in apparent signal and increase in holding current, likely indicating the loss of a stable recording configuration. We occasionally observed cases where we recorded outward currents in situations where inward currents would be expected e.g. sometimes one of two channels recording a given group of transfected cells would display inward currents while the other channel displayed outward currents. We do not have a satisfactory explanation for this phenomenon, but we excluded assays where this was observed. We also encountered apparent photovoltaic artifacts in some experiments during the highest intensity stimulation in the 440 and 475 nm channels. These artifacts were encountered consistently on a subset of the recording channels; this idiosyncrasy is likely a consequence of the exact geometry of our light path with respect to the recording electrodes and occlusions created by the patch clamp system and plates.

For a dataset to be considered for fitting, a minimum peak photocurrent response of at least 100 pA was required. For an assay to be included in the dataset of fitted parameters (i.e. considered a successful recording), the model fit needed to achieve an average R² value of at least 0.25 on holdout data in 5-fold cross validation, though typical values were much higher (Fig S3). Inspection of the fits revealed that they generally passed the ‘eye-test’ though we noted some shortcomings. Most saliently, there were cases where predictions exhibited a wavelength-dependent bias; specifically, for some more red-shifted opsins, fitted action surfaces would sometimes systematically underestimate the size of currents for 440 nm and 475 nm stimulation. This likely reflects misspecification in the first stage of the action surface model: a simple Gaussian is inadequate to capture the true shape of the action spectrum. For more red-shifted opsins, the Gaussian fails to account for the spectral “shoulder” that is seen in bluer wavelengths of action/absorption spectra for CRs. More complex models are able to account for this, e.g. a mixture of Gaussians ^26^, the Lamb equation ^138^, or a parametric Weibull function ^14^. However, we encountered apparent issues with identifiability when we attempted to use these models as the action spectrum function in lieu of a simple Gaussian, which is perhaps not surprising given our sparse sampling of spectral space. Kinetic analysis To characterize the closing kinetics of the opsin channels, the photocurrent decay following the cessation of the light stimulus was fitted with a single exponential function. This analysis was performed on sweeps that exhibited a peak current of at least 80 pA. The off-kinetics were modeled using the following equation:

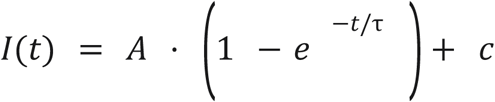

where I(t) is the current as a function of time t, *A* is the amplitude of the exponential component, τis the time constant of the exponential decay, representing the off-kinetics, and *c* is a constant representing the steady-state current offset. The fitting was performed on a 500 ms window immediately following the end of the photostimulation. For a fit to be included, it needed to achieve an R^2^ value of at least 0.7. For an assay to be considered to have reliable off-kinetics, a minimum of 10 sweeps meeting these criteria were required. The final reported off-kinetic value for an opsin variant is the median τ_off_ from all the sweeps that passed these quality control filters.

### Analysis of Manual Patch-Clamp Data

All data from manual patch-clamp recordings were processed and analyzed using custom Python scripts, using a similar methodology to the automated patch-clamp data. For each recording sweep, the raw current trace was first baseline-corrected by subtracting the median current from a defined pre-stimulus window. Both the peak current (the maximum absolute current during the light pulse) and the steady-state current (the median current in the latter part of the light pulse - typically the last 400 ms) were extracted. For each of these measurements, we fitted a Hill equation as above.

Photocurrent kinetics and desensitization were quantified from baseline-corrected current traces. The deactivation time constant (τ_off_) was determined by fitting a single exponential function to the current’s decay following light termination. As a complementary approach, a numerical time constant was also calculated by directly measuring the time required for the current to decay by 63% from its value at light offset; this was generally more robust and accounts for situations where decay is not well-described by a single exponential process, so we report these values as τ_off_ for manual patch data. The degree of desensitization during sustained illumination was assessed by calculating the ratio of the steady-state current, measured in the latter part of the illumination window, to the initial peak current.

### *In vivo* Electrophysiology Data Analysis

SpikeGLX recordings were first preprocessed with ecephys spike sorting^139^and then clustered with Kilosort 4^140^. Sorted clusters were manually curated with phy^141^. Only well-isolated single units were included in the analysis. Spike rate was calculated using 10ms bins with causal exponential decay filter with τ= 25 ms. Mean baseline firing rate in 1s prestimulus period was subtracted from traces. Z-scores were calculated using pre photostimulation spike rates and applied to photostim trials.

### Histology

Following *in vivo* recordings, mice were deeply anesthetized using ketamine and transcardially perfused with cold PBS followed by 4% paraformaldehyde (PFA). Brains were extracted, post-fixed in 4% PFA overnight at 4°C, and then cryoprotected by immersion in 30% sucrose in PBS at 4°C until sunk. Coronal cryosections (typically 50 µm thick) were collected around the injection and recording sites. Sections were stained with DAPI (4′,6-diamidino-2-phenylindole) to visualize cell nuclei, and fluorescent images were acquired to visualize DAPI and mScarlet (ChRmine and WAChR variant expression). DiI (lipophilic tracer) was applied to a subset of Neuropixels probe before insertion to subsequently visualize the probe track, however probe track can be identified in the absence of Dil staining. Images were acquired on a Zeiss Axioscan 7 and a Zeiss LSM 980.

### Protein Structure Prediction

Predicted protein structures of WiChR and WiChR F240A (“WAChR”) were generated using Boltz2^132^. Each protein was predicted as a trimer because the closely related HcKCR1 is a trimer^41^. Additionally, each protein was predicted with a single retinal cofactor using a bond constraint between the C15 position of retinal and the NZ position of the Schiff base lysine at position 251 of WiChR. Ion channel cavities were mapped out using PDBStruct HOLLOW^142^ with grid spacing set to 0.3 and internal probe setting set to 1.3. Predictions were performed on a computer equipped with 4x NVIDIA A100 GPUs, 128 vCPU, and 1 TB RAM.

### Sequence Alignment

Sequence alignment between HcKCR1, HcKCR2, and WiChR was calculated using MAFFT-linsi^110^ and visualized with ESPript3^143^.

## Acknowledgements

We would like to express our appreciation for Sabrina Thompson and Kevin Smith for histology support, to Rebecca Chowdhury for generating iPSC-derived human neurons, to Lena Bengtsson for technical assistance, to Mo Eltaeb, Silvio Ortiz, and Ben Shababo for automation assistance, to Kacylene Sangalang and Yuri Humrich for cell culture, to Ben Rakela for virus injections, to Eric Knudsen for assistance with *in vivo* electrophysiology, and to Max Hodak for providing guidance and support.

## Author Contributions

A.T. performed cell culture, automated patch-clamp, manual patch-clamp, and neuropixel experiments, analyzed data, and wrote the manuscript. A.Nava designed and performed mutagenesis experiments, wrote the manuscript, wrote the machine-learning pipeline, and analyzed data. S.M. performed cell culture, molecular biology, mutagenesis, and automated patch-clamp experiments. A.M. designed the project, provided technical direction, and wrote the manuscript. A. Naka designed the project, directed technical operations, wrote the manuscript, analyzed data, and designed and wrote the machine-learning pipeline.

## References

1. Pan, Z.-H., Ganjawala, T.H., Lu, Q., Ivanova, E., and Zhang, Z. (2014). ChR2 mutants at L132 and T159 with improved operational light sensitivity for vision restoration. PLoS One 9, e98924.

2. Gaub, B.M., Berry, M.H., Holt, A.E., Isacoff, E.Y., and Flannery, J.G. (2015). Optogenetic vision restoration using rhodopsin for enhanced sensitivity. Mol. Ther. 23, 1562–1571.

3. Gilhooley, M.J., Lindner, M., Palumaa, T., Hughes, S., Peirson, S.N., and Hankins, M.W. (2022). A systematic comparison of optogenetic approaches to visual restoration. Mol. Ther. Methods Clin. Dev. 25, 111–123.

4. Sahel, J.-A., Boulanger-Scemama, E., Pagot, C., Arleo, A., Galluppi, F., Martel, J.N., Esposti, S.D., Delaux, A., de Saint Aubert, J.-B., de Montleau, C.,collab et al. (2021). Partial recovery of visual function in a blind patient after optogenetic therapy. Nat. Med. 27, 1223–1229.

5. Klapoetke, N.C., Murata, Y., Kim, S.S., Pulver, S.R., Birdsey-Benson, A., Cho, Y.K., Morimoto, T.K., Chuong, A.S., Carpenter, E.J., Tian, Z., et al. (2014). Independent optical excitation of distinct neural populations. Nat. Methods 11, 338–346.

6. Govorunova, E.G., Gou, Y., Sineshchekov, O.A., Li, H., Lu, X., Wang, Y., Brown, L.S., St-Pierre, F., Xue, M., and Spudich, J.L. (2022). Kalium channelrhodopsins are natural light-gated potassium channels that mediate optogenetic inhibition. Nat. Neurosci. 25, 967–974.

7. Marshel, J.H., Kim, Y.S., Machado, T.A., Quirin, S., Benson, B., Kadmon, J., Raja, C., Chibukhchyan, A., Ramakrishnan, C., Inoue, M., et al. (2019). Cortical layer-specific critical dynamics triggering perception. Science. 10.1126/science.aaw5202.

8. Govorunova, E.G., Sineshchekov, O.A., Rodarte, E.M., Janz, R., Morelle, O., Melkonian, M., Wong, G.K.-S., and Spudich, J.L. (2017). The Expanding Family of Natural Anion Channelrhodopsins Reveals Large Variations in Kinetics, Conductance, and Spectral Sensitivity. Sci. Rep. 7, 43358.

9. Govorunova, E.G., Sineshchekov, O.A., Li, H., Wang, Y., Brown, L.S., Palmateer, A., Melkonian, M., Cheng, S., Carpenter, E., Patterson, J., et al. (2021). Cation and anion channelrhodopsins: Sequence motifs and taxonomic distribution. MBio 12, e0165621.

10. Govorunova, E.G., Sineshchekov, O.A., Li, H., Wang, Y., Brown, L.S., and Spudich, J.L. (2020). RubyACRs, nonalgal anion channelrhodopsins with highly red-shifted absorption. Proc. Natl. Acad. Sci. U. S. A. 117, 22833–22840.

11. Govorunova, E.G., Sineshchekov, O.A., Li, H., Gou, Y., Chen, H., Yang, S., Wang, Y., Mitchell, S., Palmateer, A., Brown, L.S., et al. (2025). Blue-shifted ancyromonad channelrhodopsins for multiplex optogenetics. bioRxivorg. 10.1101/2025.02.24.639930.

12. Govorunova, E.G., Sineshchekov, O.A., and Spudich, J.L. (2021). Emerging Diversity of Channelrhodopsins and Their Structure-Function Relationships. Front. Cell. Neurosci. 15, 800313.

13. Tanaka, T., Hososhima, S., Yamashita, Y., Sugimoto, T., Nakamura, T., Shigemura, S., Iida, W., Sano, F.K., Oda, K., Uchihashi, T., et al. (2024). The high-light-sensitivity mechanism and optogenetic properties of the bacteriorhodopsin-like channelrhodopsin GtCCR4. Mol. Cell 0. 10.1016/j.molcel.2024.08.016.

14. Oda, K., Vierock, J., Oishi, S., Taniguchi, R., Yamashita, K., Nishizawa, T., Hegemann, P., and Nureki, O. (2018). Crystal structure of the red light-activated channelrhodopsin Chrimson. Preprint at Worldwide Protein Data Bank, 10.2210/pdb5zih/pdb.

15. Bedbrook, C.N., Yang, K.K., Robinson, J.E., Mackey, E.D., Gradinaru, V., and Arnold, F.H. (2019). Machine learning-guided channelrhodopsin engineering enables minimally invasive optogenetics. Nat. Methods 16, 1176–1184.

16. Bedbrook, C.N., Yang, K.K., Rice, A.J., Gradinaru, V., and Arnold, F.H. (2017). Machine learning to design integral membrane channelrhodopsins for efficient eukaryotic expression and plasma membrane localization. PLoS Comput. Biol. 13, e1005786.

17. Gradinaru, V., Zhang, F., Ramakrishnan, C., Mattis, J., Prakash, R., Diester, I., Goshen, I., Thompson, K.R., and Deisseroth, K. (2010). Molecular and cellular approaches for diversifying and extending optogenetics. Cell 141, 154–165.

18. Kato, H.E., Kim, Y.S., Paggi, J.M., Evans, K.E., Allen, W.E., Richardson, C., Inoue, K., Ito, S., Ramakrishnan, C., Fenno, L.E., et al. (2018). Structural mechanisms of selectivity and gating in anion channelrhodopsins. Nature 561, 349–354.

19. Lin, J.Y., Knutsen, P.M., Muller, A., Kleinfeld, D., and Tsien, R.Y. (2013). ReaChR: a red-shifted variant of channelrhodopsin enables deep transcranial optogenetic excitation. Nat. Neurosci. 16, 1499–1508.

20. Berndt, A., Yizhar, O., Gunaydin, L.A., Hegemann, P., and Deisseroth, K. (2009). Bi-stable neural state switches. Nat. Neurosci. 12, 229–234.

21. Yizhar, O., Fenno, L.E., Prigge, M., Schneider, F., Davidson, T.J., O’Shea, D.J., Sohal, V.S., Goshen, I., Finkelstein, J., Paz, J.T., et al. (2011). Neocortical excitation/inhibition balance in information processing and social dysfunction. Nature 477, 171–178.

22. Gunaydin, L.A., Yizhar, O., Berndt, A., Sohal, V.S., Deisseroth, K., and Hegemann, P. (2010). Ultrafast optogenetic control. Nat. Neurosci. 13, 387–392.

23. Berndt, A., Schoenenberger, P., Mattis, J., Tye, K.M., Deisseroth, K., Hegemann, P., and Oertner, T.G. (2011). High-efficiency channelrhodopsins for fast neuronal stimulation at low light levels. Proc. Natl. Acad. Sci. U. S. A. 108, 7595–7600.

24. Prigge, M., Schneider, F., Tsunoda, S.P., Shilyansky, C., Wietek, J., Deisseroth, K., and Hegemann, P. (2012). Color-tuned channelrhodopsins for multiwavelength optogenetics. J. Biol. Chem. 287, 31804–31812.

25. Oppermann, J., Rozenberg, A., Fabrin, T., GonzalezCabrera, C., Béjà, O., Prigge, M., and Hegemann, P. (2023). Robust Optogenetic Inhibition with Red-light-sensitive Anion-conducting Channelrhodopsins. bioRxiv, 2023.06.09.544329. 10.1101/2023.06.09.544329.

26. Kato, H.E., Kamiya, M., Sugo, S., Ito, J., Taniguchi, R., Orito, A., Hirata, K., Inutsuka, A., Yamanaka, A., Maturana, A.D., et al. (2015). Atomistic design of microbial opsin-based blue-shifted optogenetics tools. Nat. Commun. 6, 7177.

27. Too, L.K., Shen, W., Protti, D.A., Sawatari, A., A Black, D., Leamey, C.A., Y Huang, J., Lee, S.-R., E Mathai, A., Lisowski, L., et al. (2022). Optogenetic restoration of high sensitivity vision with bReaChES, a red-shifted channelrhodopsin. Sci. Rep. 12, 19312.

28. Mardinly, A.R., Oldenburg, I.A., Pégard, N.C., Sridharan, S., Lyall, E.H., Chesnov, K., Brohawn, S.G., Waller, L., and Adesnik, H. (2018). Precise multimodal optical control of neural ensemble activity. Nat. Neurosci. 21, 881–893.

29. Cho, Y.K., Park, D., Yang, A., Chen, F., Chuong, A.S., Klapoetke, N.C., and Boyden, E.S. (2019). Multidimensional screening yields channelrhodopsin variants having improved photocurrent and order-of-magnitude reductions in calcium and proton currents. J. Biol. Chem. 294, 3806–3821.

30. Fernandez Lahore, R.G., Pampaloni, N.P., Peter, E., Heim, M.-M., Tillert, L., Vierock, J., Oppermann, J., Walther, J., Schmitz, D., Owald, D., et al. (2022). Calcium-permeable channelrhodopsins for the photocontrol of calcium signalling. Nat. Commun. 13, 7844.

31. Vierock, J., Grimm, C., Nitzan, N., and Hegemann, P. (2017). Molecular determinants of proton selectivity and gating in the red-light activated channelrhodopsin Chrimson. Sci. Rep. 7, 9928.

32. Bi, X., Beck, C., and Gong, Y. (2022). A kinetic-optimized CoChR variant with enhanced high-frequency spiking fidelity. Biophys. J. 121, 4166–4178.

33. Ganjawala, T.H., Lu, Q., Fenner, M.D., Abrams, G.W., and Pan, Z.-H. (2019). Improved CoChR variants restore visual acuity and contrast sensitivity in a mouse model of blindness under ambient light conditions. Mol. Ther. 27, 1195–1205.

34. Watanabe, Y., Sugano, E., Tabata, K., Hatakeyama, A., Sakajiri, T., Fukuda, T., Ozaki, T., Suzuki, T., Sayama, T., and Tomita, H. (2021). Development of an optogenetic gene sensitive to daylight and its implications in vision restoration. NPJ Regen. Med. 6, 64.

35. Rodriguez-Rozada, S., Wietek, J., Tenedini, F., Sauter, K., Dhiman, N., Hegemann, P., Soba, P., and Wiegert, J.S. (2022). Aion is a bistable anion-conducting channelrhodopsin that provides temporally extended and reversible neuronal silencing. Commun Biol 5, 687.

36. Wietek, J., Rodriguez-Rozada, S., Tutas, J., Tenedini, F., Grimm, C., Oertner, T.G., Soba, P., Hegemann, P., and Wiegert, J.S. (2017). Anion-conducting channelrhodopsins with tuned spectra and modified kinetics engineered for optogenetic manipulation of behavior. Sci. Rep. 7, 14957.

37. Rodriguez-Rozada, S., Wietek, J., Tenedini, F., Sauter, K., Hegemann, P., Soba, P., and Simon Wiegert, J. (2022). Temporally extended and reversible neuronal silencing with Aion. bioRxiv, 2022.02.25.481932. 10.1101/2022.02.25.481932.

38. Wietek, J., and Prigge, M. (2016). Enhancing Channelrhodopsins: An Overview. In Optogenetics: Methods and Protocols, A. Kianianmomeni, ed. (Springer New York), pp. 141–165.

39. Vierock, J., Rodriguez-Rozada, S., Dieter, A., Pieper, F., Sims, R., Tenedini, F., Bergs, A.C.F., Bendifallah, I., Zhou, F., Zeitzschel, N., et al. (2021). BiPOLES is an optogenetic tool developed for bidirectional dual-color control of neurons. Nat. Commun. 12, 4527.

40. Kishi, K.E., Kim, Y.S., Fukuda, M., Inoue, M., Kusakizako, T., Wang, P.Y., Ramakrishnan, C., Byrne, E.F.X., Thadhani, E., Paggi, J.M., et al. (2022). Structural basis for channel conduction in the pump-like channelrhodopsin ChRmine. Cell 185, 672–689.e23.

41. Tajima, S., Kim, Y.S., Fukuda, M., Jo, Y., Wang, P.Y., Paggi, J.M., Inoue, M., Byrne, E.F.X., Kishi, K.E., Nakamura, S., et al. (2023). Structural basis for ion selectivity in potassium-selective channelrhodopsins. Cell 186, 4325–4344.e26.

42. Kleinlogel, S., Feldbauer, K., Dempski, R.E., Fotis, H., Wood, P.G., Bamann, C., and Bamberg, E. (2011). Ultra light-sensitive and fast neuronal activation with the Ca^2^+-permeable channelrhodopsin CatCh. Nat. Neurosci. 14, 513–518.

43. Dawydow, A., Gueta, R., Ljaschenko, D., Ullrich, S., Hermann, M., Ehmann, N., Gao, S., Fiala, A., Langenhan, T., Nagel, G., et al. (2014). Channelrhodopsin-2-XXL, a powerful optogenetic tool for low-light applications. Proc. Natl. Acad. Sci. U. S. A. 111, 13972–13977.

44. Sridharan, S., Gajowa, M.A., Ogando, M.B., Jagadisan, U.K., Abdeladim, L., Sadahiro, M., Bounds, H.A., Hendricks, W.D., Turney, T.S., Tayler, I., et al. (2022). High-performance microbial opsins for spatially and temporally precise perturbations of large neuronal networks. Neuron 110, 1139–1155.e6.

45. Alekseev, A., Hunniford, V., Zerche, M., Jeschke, M., El May, F., Vavakou, A., Siegenthaler, D., Hüser, M.A., Kiehn, S.M., Garrido-Charles, A., et al. (2025). Efficient and sustained optogenetic control of sensory and cardiac systems. Nat. Biomed. Eng. 10.1038/s41551-025-01461-1.

46. Baker, C.A., Elyada, Y.M., Parra, A., and Bolton, M.M. (2016). Cellular resolution circuit mapping with temporal-focused excitation of soma-targeted channelrhodopsin. Elife 5, e14193.

47. Messier, J.E., Chen, H., Cai, Z.-L., and Xue, M. (2018). Targeting light-gated chloride channels to neuronal somatodendritic domain reduces their excitatory effect in the axon. Elife 7. 10.7554/eLife.38506.

48. Shemesh, O.A., Tanese, D., Zampini, V., Linghu, C., Piatkevich, K., Ronzitti, E., Papagiakoumou, E., Boyden, E.S., and Emiliani, V. (2017). Temporally precise single-cell-resolution optogenetics. Nat. Neurosci. 20, 1796–1806.

49. Gong, X., Mendoza-Halliday, D., Ting, J.T., Kaiser, T., Sun, X., Bastos, A.M., Wimmer, R.D., Guo, B., Chen, Q., Zhou, Y., et al. (2020). An Ultra-Sensitive Step-Function Opsin for Minimally Invasive Optogenetic Stimulation in Mice and Macaques. Neuron 107, 38–51.e8.

50. Vierock, J., Shiewer, E., Grimm, C., Rozenberg, A., Chen, I.-W., Tillert, L., Castro Scalise, A.G., Casini, M., Augustin, S., Tanese, D., et al. (2022). WiChR, a highly potassium-selective channelrhodopsin for low-light one- and two-photon inhibition of excitable cells. Sci. Adv. 8, eadd7729.

51. Sineshchekov, O.A., Govorunova, E.G., Li, H., and Spudich, J.L. (2017). Bacteriorhodopsin-like channelrhodopsins: Alternative mechanism for control of cation conductance. Proc. Natl. Acad. Sci. U. S. A. 114, E9512–E9519.

52. Parnami, K., and Bhattacharyya, A. (2023). Current approaches to vision restoration using optogenetic therapy. Front. Cell. Neurosci. 17, 1236826.

53. Poboży, K., Poboży, T., Domański, P., Derczyński, M., Konarski, W., and Domańska-Poboża, J. (2025). Evolution of light-sensitive proteins in optogenetic approaches for vision restoration: A comprehensive review. Biomedicines 13, 429.

54. Tucker, K., Sridharan, S., Adesnik, H., and Brohawn, S.G. (2022). Cryo-EM structures of the channelrhodopsin ChRmine in lipid nanodiscs. Nat. Commun. 13, 4842.

55. Gradinaru, V., Thompson, K.R., and Deisseroth, K. (2008). eNpHR: a Natronomonas halorhodopsin enhanced for optogenetic applications. Brain Cell Biol. 36, 129–139.

56. Zerche, M., Wrobel, C., Kusch, K., Moser, T., and Mager, T. (2023). Channelrhodopsin fluorescent tag replacement for clinical translation of optogenetic hearing restoration. Mol Ther Methods Clin Dev 29, 202–212.

57. Keppeler, D., Merino, R.M., Lopez de la Morena, D., Bali, B., Huet, A.T., Gehrt, A., Wrobel, C., Subramanian, S., Dombrowski, T., Wolf, F., et al. (2018). Ultrafast optogenetic stimulation of the auditory pathway by targeting-optimized Chronos. EMBO J. 37. 10.15252/embj.201899649.

58. Shepard, B.D., Natarajan, N., Protzko, R.J., Acres, O.W., and Pluznick, J.L. (2013). A cleavable N-terminal signal peptide promotes widespread olfactory receptor surface expression in HEK293T cells. PLoS One 8, e68758.

59. Yang, S. (2022). Characterization and engineering of photoreceptors with improved properties for optogenetic application.

60. Chen, F., Yu, Y., and Shen, Y. (2020). A high sensitivity and kinetics of channelrhodopsin PsCatCh2.0 restores vision function in retinitis pigmentosa mouse model. Investigative Ophthalmology & Visual Science 61, 2733–2733.

61. Govorunova, E.G., Sineshchekov, O.A., Brown, L.S., and Spudich, J.L. (2022). Biophysical characterization of light-gated ion channels using planar automated patch clamp. Front. Mol. Neurosci. 15, 976910.

62. Yamauchi, Y., Konno, M., Ito, S., Tsunoda, S.P., Inoue, K., and Kandori, H. (2017). Molecular properties of a DTD channelrhodopsin from Guillardia theta. Biophys. Physicobiol. 14, 57–66.

63. Morizumi, T., Kim, K., Li, H., Govorunova, E.G., Sineshchekov, O.A., Wang, Y., Zheng, L., Bertalan, É., Bondar, A.-N., Askari, A., et al. (2023). Structures of channelrhodopsin paralogs in peptidiscs explain their contrasting K+ and Na+ selectivities. Nat. Commun. 14, 4365.

64. Yang, K.K., Wu, Z., and Arnold, F.H. (2019). Machine-learning-guided directed evolution for protein engineering. Nat. Methods 16, 687–694.

65. Mattis, J., Tye, K.M., Ferenczi, E.A., Ramakrishnan, C., O’Shea, D.J., Prakash, R., Gunaydin, L.A., Hyun, M., Fenno, L.E., Gradinaru, V., et al. (2011). Principles for applying optogenetic tools derived from direct comparative analysis of microbial opsins. Nat. Methods 9, 159–172.

66. Mager, T., Lopez de la Morena, D., Senn, V., Schlotte, J., D Errico, A., Feldbauer, K., Wrobel, C., Jung, S., Bodensiek, K., Rankovic, V., et al. (2018). High frequency neural spiking and auditory signaling by ultrafast red-shifted optogenetics. Nat. Commun. 9, 1750.

67. Fong, V.C., Le, B.M., Stefanov, A., Lee, V., Park, S., Sivakumar, A., Spatny, S., Visel, M., Taylor, W.R., Brohawn, S.G., et al. (2025). Optogenetic restoration of high-sensitivity vision using ChRmine- and ChroME-based channelrhodopsins. Sci. Rep. 15. 10.1038/s41598-025-04286-9.

68. Lu, Q.I., Wright, A., Abrams, G.W., and Pan, Z.-H. (2023). AAV dose-dependent transduction efficiency in RGCs and functional efficacy of CoChR3M-mediated vision restoration. Invest. Ophthalmol. Vis. Sci. 64, 214–214.

69. Berndt, A., Lee, S.Y., Wietek, J., Ramakrishnan, C., Steinberg, E.E., Rashid, A.J., Kim, H., Park, S., Santoro, A., Frankland, P.W., et al. (2016). Structural foundations of optogenetics: Determinants of channelrhodopsin ion selectivity. Proc. Natl. Acad. Sci. U. S. A. 113, 822–829.

70. Morizumi, T., Kim, K., Li, H., Nag, P., Dogon, T., Sineshchekov, O.A., Wang, Y., Brown, L.S., Hwang, S., Sun, H., et al. (2025). Structural insights into light-gating of potassium-selective channelrhodopsin. Nat. Commun. 16, 1283.

71. Sineshchekov, O.A., Govorunova, E.G., Li, H., Wang, Y., and Spudich, J.L. (2023). Channel Gating in Kalium Channelrhodopsin Slow Mutants. J. Mol. Biol., 168298.

72. Zhang, M., Shan, Y., Zhao, L., Li, X., and Pei, D. (2023). Ion selectivity and activation mechanism for kalium channelrhodopsins. bioRxiv, 2023.07.22.550149. 10.1101/2023.07.22.550149.

73. Shan, Y., Zhao, L., Chen, M., Li, X., Zhang, M., and Pei, D. (2024). Channelrhodopsins with distinct chromophores and binding patterns. Nat. Commun. 15, 1–10.

74. Govorunova, E.G., Sineshchekov, O.A., Janz, R., Liu, X., and Spudich, J.L. (2015). NEUROSCIENCE. Natural light-gated anion channels: A family of microbial rhodopsins for advanced optogenetics. Science 349, 647–650.

75. Hososhima, S., Ueno, S., Okado, S., Inoue, K.-I., Konno, M., Yamauchi, Y., Inoue, K., Terasaki, H., Kandori, H., and Tsunoda, S.P. (2023). A light-gated cation channel with high reactivity to weak light. Sci. Rep. 13, 7625.

76. Govorunova, E.G., Sineshchekov, O.A., Li, H., Janz, R., and Spudich, J.L. (2013). Characterization of a highly efficient blue-shifted channelrhodopsin from the marine alga Platymonas subcordiformis. J. Biol. Chem. 288, 29911–29922.

77. Oppermann, J., Fischer, P., Silapetere, A., Liepe, B., Rodriguez-Rozada, S., Flores-Uribe, J., Peter, E., Keidel, A., Vierock, J., Kaufmann, J., et al. (2019). MerMAIDs: a family of metagenomically discovered marine anion-conducting and intensely desensitizing channelrhodopsins. Nat. Commun. 10, 3315.

78. Labudda, K., Norahan, M.J., Hübner, L.-M., Althoff, P., Gerwert, K., Lübben, M., Rudack, T., and Kötting, C. (2025). A second photoactivatable state of the anion-conducting channelrhodopsin GtACR1 empowers persistent activity. Commun. Biol. 8, 1183.

79. Kuhne, J., Vierock, J., Tennigkeit, S.A., Dreier, M.-A., Wietek, J., Petersen, D., Gavriljuk, K., El-Mashtoly, S.F., Hegemann, P., and Gerwert, K. (2019). Unifying photocycle model for light adaptation and temporal evolution of cation conductance in channelrhodopsin-2. Proc. Natl. Acad. Sci. U. S. A. 116, 9380–9389.

80. Duan, X., Zhang, C., Wu, Y., Ju, J., Xu, Z., Li, X., Liu, Y., Ohdah, S., Constantin, O.M., Pan, Y., et al. (2025). Suppression of epileptic seizures by transcranial activation of K+-selective channelrhodopsin. Nat. Commun. 16, 559.

81. Duan, X., Zhang, C., Ott, S., Zhang, Z., Ruse, C., Dannhaeuser, S., Jacobi, R., Ehmann, N., Sachidanandan, D., Nagel, G., et al. (2025). Stabilized ion selectivity corrects activation drift in kalium channelrhodopsins. bioRxiv. 10.1101/2025.05.20.654571.

82. Knudsen, E.B., Zappitelli, K., Brown, J., Reeder, J., Smith, K.S., Rostov, M., Choi, J., Rochford, A., Slager, N., Miura, S.K., et al. (2023). A thin-film optogenetic visual prosthesis. bioRxiv, 2023.01.31.526482. 10.1101/2023.01.31.526482.

83. Drew, L. (2025). Restoring vision with optogenetics. Nature 639, S7–S9.

84. Bindels, D.S., Haarbosch, L., van Weeren, L., Postma, M., Wiese, K.E., Mastop, M., Aumonier, S., Gotthard, G., Royant, A., Hink, M.A., et al. (2017). mScarlet: a bright monomeric red fluorescent protein for cellular imaging. Nat. Methods 14, 53–56.

85. Zulkower, V., and Rosser, S. (2020). DNA Chisel, a versatile sequence optimizer. Bioinformatics 36, 4508–4509.

86. Schmidt, M., Lee, N., Zhan, C., Roberts, J.B., Nava, A.A., Keiser, L.S., Vilchez, A.A., Chen, Y., Petzold, C.J., Haushalter, R.W., et al. (2023). Maximizing heterologous expression of engineered type I polyketide synthases: Investigating Codon optimization strategies. ACS Synth. Biol. 12, 3366–3380.

87. Tang, L., Gao, H., Zhu, X., Wang, X., Zhou, M., and Jiang, R. (2012). Construction of “small-intelligent” focused mutagenesis libraries using well-designed combinatorial degenerate primers. Biotechniques 52, 149–158.

88. Hanssen, F., Garcia, M.U., Folkersen, L., Pedersen, A.S., Lescai, F., Jodoin, S., Miller, E., Seybold, M., Wacker, O., Smith, N., et al. (2024). Scalable and efficient DNA sequencing analysis on different compute infrastructures aiding variant discovery. NAR Genom. Bioinform. 6, lqae031.

89. Garcia, M., Juhos, S., Larsson, M., Olason, P.I., Martin, M., Eisfeldt, J., DiLorenzo, S., Sandgren, J., Díaz De Ståhl, T., Ewels, P., et al. (2020). Sarek: A portable workflow for whole-genome sequencing analysis of germline and somatic variants. F1000Res. *9*, 63.

90. Ewels, P.A., Peltzer, A., Fillinger, S., Patel, H., Alneberg, J., Wilm, A., Garcia, M.U., Di Tommaso, P., and Nahnsen, S. (2020). The nf-core framework for community-curated bioinformatics pipelines. Nat. Biotechnol. 38, 276–278.

91. Zheng, W., Wuyun, Q., Li, Y., Zhang, C., Freddolino, L., and Zhang, Y. (2024). Improving deep learning protein monomer and complex structure prediction using DeepMSA2 with huge metagenomics data. Nat. Methods 21, 279–289.

92. Mirdita, M., von den Driesch, L., Galiez, C., Martin, M.J., Söding, J., and Steinegger, M. (2017). Uniclust databases of clustered and deeply annotated protein sequences and alignments. Nucleic Acids Res. 45, D170–D176.

93. Steinegger, M., Meier, M., Mirdita, M., Vöhringer, H., Haunsberger, S.J., and Söding, J. (2019). HH-suite3 for fast remote homology detection and deep protein annotation. BMC Bioinformatics 20, 473.

94. Eddy, S.R. (2011). Accelerated profile HMM searches. PLoS Comput. Biol. 7, e1002195.

95. Nayfach, S., Páez-Espino, D., Call, L., Low, S.J., Sberro, H., Ivanova, N.N., Proal, A.D., Fischbach, M.A., Bhatt, A.S., Hugenholtz, P., et al. (2021). Metagenomic compendium of 189,680 DNA viruses from the human gut microbiome. Nat. Microbiol. *6*, 960–970.

96. . Camarillo-Guerrero, L.F., Almeida, A., Rangel-Pineros, G., Finn, R.D., and Lawley, T.D. (2021). Massive expansion of human gut bacteriophage diversity. Cell 184, 1098–1109.e9.

97. Ma, B., Lu, C., Wang, Y., Yu, J., Zhao, K., Xue, R., Ren, H., Lv, X., Pan, R., Zhang, J., et al. (2023). A genomic catalogue of soil microbiomes boosts mining of biodiversity and genetic resources. Nat. Commun. *14*, 7318.

98. Chen, I.-M.A., Chu, K., Palaniappan, K., Pillay, M., Ratner, A., Huang, J., Huntemann, M., Varghese, N., White, J.R., Seshadri, R., et al. (2019). IMG/M v.5.0: an integrated data management and comparative analysis system for microbial genomes and microbiomes. Nucleic Acids Res. 47, D666–D677.

99. Camargo, A.P., Nayfach, S., Chen, I.-M.A., Palaniappan, K., Ratner, A., Chu, K., Ritter, S.J., Reddy, T.B.K., Mukherjee, S., Schulz, F., et al. (2023). IMG/VR v4: an expanded database of uncultivated virus genomes within a framework of extensive functional, taxonomic, and ecological metadata. Nucleic Acids Res. 51, D733–D743.

100. Steinegger, M., Mirdita, M., and Söding, J. (2019). Protein-level assembly increases protein sequence recovery from metagenomic samples manyfold. Nat. Methods *16*, 603–606.

101. Suzek, B.E., Wang, Y., Huang, H., McGarvey, P.B., Wu, C.H., and UniProt Consortium (2015). UniRef clusters: a comprehensive and scalable alternative for improving sequence similarity searches. Bioinformatics 31, 926–932.

102. Steinegger, M., and Söding, J. (2018). Clustering huge protein sequence sets in linear time. Nat. Commun. 9, 2542.

103. Richardson, L., Allen, B., Baldi, G., Beracochea, M., Bileschi, M.L., Burdett, T., Burgin, J., Caballero-Pérez, J., Cochrane, G., Colwell, L.J., et al. (2023). MGnify: the microbiome sequence data analysis resource in 2023. Nucleic Acids Res. 51, D753–D759.

104. Hu, E.J., Shen, Y., Wallis, P., Allen-Zhu, Z., Li, Y., Wang, S., Wang, L., and Chen, W. (2021). LoRA: Low-Rank Adaptation of large language models. arXiv [cs.CL]. 10.48550/ARXIV.2106.09685.

105. Lin, Z., Akin, H., Rao, R., Hie, B., Zhu, Z., Lu, W., Smetanin, N., Verkuil, R., Kabeli, O., Shmueli, Y., et al. (2023). Evolutionary-scale prediction of atomic-level protein structure with a language model. Science 379, 1123–1130.

106. Ouyang-Zhang, J., Gong, C., Zhao, Y., Krähenbühl, P., Klivans, A.R., and Diaz, D.J. (2024). Distilling structural representations into protein sequence models. bioRxiv. 10.1101/2024.11.08.622579.

107. Yang, K.K., Fusi, N., and Lu, A.X. (2024). Convolutions are competitive with transformers for protein sequence pretraining. Cell Systems. 10.1016/j.cels.2024.01.008.

108. Gordon, C., Lu, A.X., and Abbeel, P. (2024). Protein language model fitness is a matter of preference. bioRxiv, 2024.10.03.616542. 10.1101/2024.10.03.616542.

109. Meier, J., Rao, R., Verkuil, R., Liu, J., Sercu, T., and Rives, A. (2021). Language models enable zero-shot prediction of the effects of mutations on protein function. bioRxiv, 2021.07.09.450648. 10.1101/2021.07.09.450648.

110. Katoh, K., and Standley, D.M. (2013). MAFFT multiple sequence alignment software version 7: improvements in performance and usability. Mol. Biol. Evol. 30, 772–780.

111. Sugihara, Y., Kourelis, J., Contreras, M.P., Pai, H., Harant, A., Selvaraj, M., Toghani, A., Martínez-Anaya, C., and Kamoun, S. (2025). Helper NLR immune protein NRC3 evolved to evade inhibition by a cyst nematode virulence effector. PLoS Genet. 21, e1011653.

112. Minh, B.Q., Schmidt, H.A., Chernomor, O., Schrempf, D., Woodhams, M.D., von Haeseler, A., and Lanfear, R. (2020). IQ-TREE 2: New models and efficient methods for phylogenetic inference in the genomic era. Mol. Biol. Evol. 37, 1530–1534.

113. Oliva, A., Pulicani, S., Lefort, V., Bréhélin, L., Gascuel, O., and Guindon, S. (2019). Accounting for ambiguity in ancestral sequence reconstruction. Bioinformatics 35, 4290–4297.

114. Eick, G.N., Bridgham, J.T., Anderson, D.P., Harms, M.J., and Thornton, J.W. (2017). Robustness of reconstructed ancestral protein functions to statistical uncertainty. Mol. Biol. Evol. 34, 247–261.

115. Eick, G.N., Colucci, J.K., Harms, M.J., Ortlund, E.A., and Thornton, J.W. (2012). Evolution of minimal specificity and promiscuity in steroid hormone receptors. PLoS Genet. 8, e1003072.

116. Bedbrook, C.N., Rice, A.J., Yang, K.K., Ding, X., Chen, S., LeProust, E.M., Gradinaru, V., and Arnold, F.H. (2017). Structure-guided SCHEMA recombination generates diverse chimeric channelrhodopsins. Proc. Natl. Acad. Sci. U. S. A. 114, E2624–E2633.

117. Wlodek, A., Kendrew, S.G., Coates, N.J., Hold, A., Pogwizd, J., Rudder, S., Sheehan, L.S., Higginbotham, S.J., Stanley-Smith, A.E., Warneck, T., et al. (2017). Diversity oriented biosynthesis via accelerated evolution of modular gene clusters. Nat. Commun. 8, 1206.

118. Dauparas, J., Anishchenko, I., Bennett, N., Bai, H., Ragotte, R.J., Milles, L.F., Wicky, B.I.M., Courbet, A., de Haas, R.J., Bethel, N., et al. (2022). Robust deep learning-based protein sequence design using ProteinMPNN. Science 378, 49–56.

119. Dauparas, J., Lee, G.R., Pecoraro, R., An, L., Anishchenko, I., Glasscock, C., and Baker, D. (2025). Atomic context-conditioned protein sequence design using LigandMPNN. Nat. Methods *22*, 717–723.

120. Emami, P., Perreault, A., Law, J., Biagioni, D., and St. John, P. (2023). Plug & play directed evolution of proteins with gradient-based discrete MCMC. Mach. Learn. Sci. Technol. 4, 025014.

121. Hawkins-Hooker, A., Surana, S., Simons, J., Kmec, J., Bent, O., and Duckworth, P. (2024). Likelihood-based Fine-tuning of Protein Language Models for Few-shot Fitness Prediction and Design. Bioinformatics.

122. Rohanian, O., Nouriborji, M., Kouchaki, S., and Clifton, D.A. (2023). On the effectiveness of compact biomedical transformers. Bioinformatics 39, btad103.

123. He, K., Fan, H., Wu, Y., Xie, S., and Girshick, R. (2020). Momentum Contrast for Unsupervised Visual Representation Learning. In Proceedings of the IEEE/CVF Conference on Computer Vision and Pattern Recognition, pp. 9729–9738.

124. Chen, Z., Badrinarayanan, V., Lee, C.-Y., and Rabinovich, A. (2017). GradNorm: Gradient normalization for adaptive loss balancing in deep multitask networks. arXiv [cs.CV].

125. Yu, T., Kumar, S., Gupta, A., Levine, S., Hausman, K., and Finn, C. (2020). Gradient Surgery for Multi-Task Learning. arXiv [cs.LG].

126. . Johnson, S.R., Fu, X., Viknander, S., Goldin, C., Monaco, S., Zelezniak, A., and Yang, K.K. (2025). Computational scoring and experimental evaluation of enzymes generated by neural networks. Nat. Biotechnol. 43, 396–405.

127. Damani, F., Brookes, D.H., Sternlieb, T., Webster, C., Malina, S., Jajoo, R., Lin, K., and Sinai, S. (2023). Beyond the training set: an intuitive method for detecting distribution shift in model-based optimization. arXiv [cs.LG]. 10.48550/ARXIV.2311.05363.

128. Bernhofer, M., and Rost, B. (2022). TMbed: transmembrane proteins predicted through language model embeddings. BMC Bioinformatics 23, 326.

129. Jumper, J., Evans, R., Pritzel, A., Green, T., Figurnov, M., Ronneberger, O., Tunyasuvunakool, K., Bates, R., Žídek, A., Potapenko, A., et al. (2021). Highly accurate protein structure prediction with AlphaFold. Nature 596, 583–589.

130. Evans, R., O’Neill, M., Pritzel, A., Antropova, N., Senior, A., Green, T., Žídek, A., Bates, R., Blackwell, S., Yim, J., et al. (2021). Protein complex prediction with AlphaFold-Multimer. Bioinformatics.

131. Mirdita, M., Schütze, K., Moriwaki, Y., Heo, L., Ovchinnikov, S., and Steinegger, M. (2022). ColabFold: making protein folding accessible to all. Nat. Methods *19*, 679–682.

132. Passaro, S., Corso, G., Wohlwend, J., Reveiz, M., Thaler, S., Somnath, V.R., Getz, N., Portnoi, T., Roy, J., Stark, H., et al. (2025). Boltz-2: Towards accurate and efficient binding affinity prediction. bioRxivorg. 10.1101/2025.06.14.659707.

133. Alamdari, S., Thakkar, N., van den Berg, R., Lu, A.X., Fusi, N., Amini, A.P., and Yang, K.K. (2023). Protein generation with evolutionary diffusion: sequence is all you need. bioRxiv, 2023.09.11.556673. 10.1101/2023.09.11.556673.

134. Auer, P. (2003). Using confidence bounds for exploitation-exploration trade-offs. J. Mach. Learn. Res. *3*, 397–422.

135. Daulton, S., Balandat, M., and Bakshy, E. (2021). Parallel Bayesian optimization of multiple noisy objectives with expected hypervolume improvement. arXiv [cs.LG]. 10.48550/ARXIV.2105.08195.

136. Telezhkin, V., Schnell, C., Yarova, P., Yung, S., Cope, E., Hughes, A., Thompson, B.A., Sanders, P., Geater, C., Hancock, J.M., et al. (2016). Forced cell cycle exit and modulation of GABAA, CREB, and GSK3β signaling promote functional maturation of induced pluripotent stem cell-derived neurons. Am. J. Physiol. Cell Physiol. 310, C520–C541.

137. Lu, Q., Wright, A., and Pan, Z.-H. (2024). AAV dose-dependent transduction efficiency in retinal ganglion cells and functional efficacy of optogenetic vision restoration. Gene Ther. 31, 572–579.

138. Govardovskii, V.I., Fyhrquist, N., Reuter, T., Kuzmin, D.G., and Donner, K. (2000). In search of the visual pigment template. Vis. Neurosci. *17*, 509–528.

139. Colonell, J. ecephys_spike_sorting: Modules for processing extracellular electrophysiology data from Neuropixels probes (Github).

140. Pennington, J., Pachitariu, M., Rossant, C., Steinmetz, N., carsen-stringer, Sridhar, S., Colonell, J., Bondy, A., Winter, O., Ressmeyer, R., et al. (2025). MouseLand/Kilosort: Kilosort v4.1.1 (Zenodo) 10.5281/ZENODO.3597474.

141. Rossant, C., Harris, K., Carandini, M., and International Brain Laboratory phy: interactive visualization and manual spike sorting of large-scale ephys data. Phy. https://github.com/cortex-lab/phy.

142. Ho, B.K., and Gruswitz, F. (2008). HOLLOW: generating accurate representations of channel and interior surfaces in molecular structures. BMC Struct. Biol. 8, 49.

143. Robert, X., and Gouet, P. (2014). Deciphering key features in protein structures with the new ENDscript server. Nucleic Acids Res. 42, W320–W324.

