## Supplementary figures and tables for "WAChRs are excitatory opsins sensitive to indoor lighting"

### Supplementary Figure 1

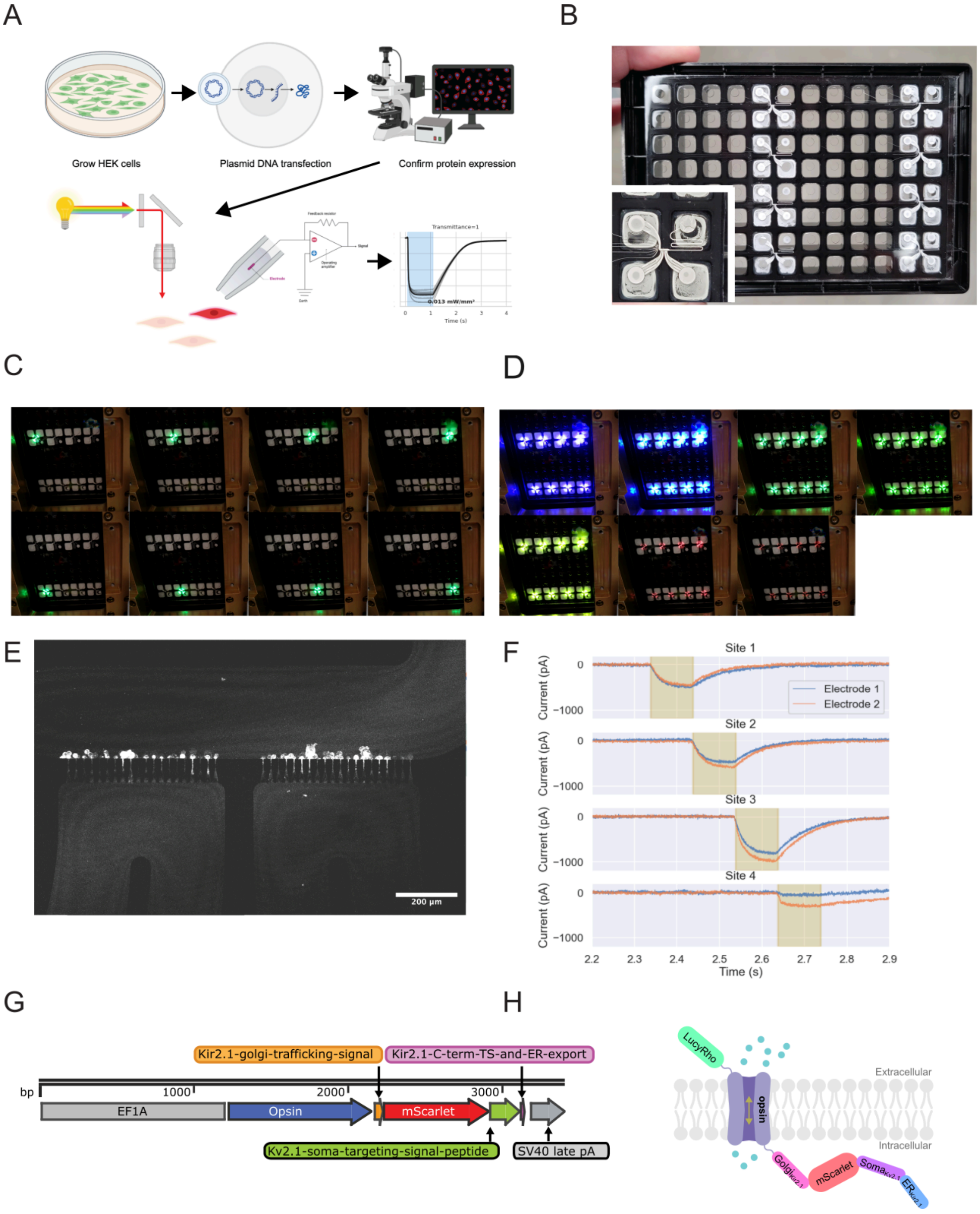


**Supplementary Figure 1.** Details of the autopatch assay
**(A)** Cartoon illustrating design of functional opsin screen
**(B)** A microfluidic autopatch plate filled with white ink to visualize electrode recording sites and serve as a calibration for the light path.
(**C**) A series of images showing point-by-point illumination of each electrode site using a single wavelength of light.

**(D)** Mean images showing how each electrode site is illuminated on each trial using 7 different wavelengths of light.
(**E**) Confocal microscope showing mScarlet-expressing HEK293T cells attached to the recording sites of the microfluidic patch plate
(**F**) Traces of ChRmine-expressing HEK293T cell showing both ensemble electrodes at each recording site as the galvo mirror targets light to the subsequent recording site.
(**G**) Linearized DNA map of our opsin-expression cassette
(**H**) Cartoon showing location of expression tags relative to plasma membrane. The LucyRho tag is only included on certain constructs as a means to improve expression and trafficking.

### Supplementary Figure 2

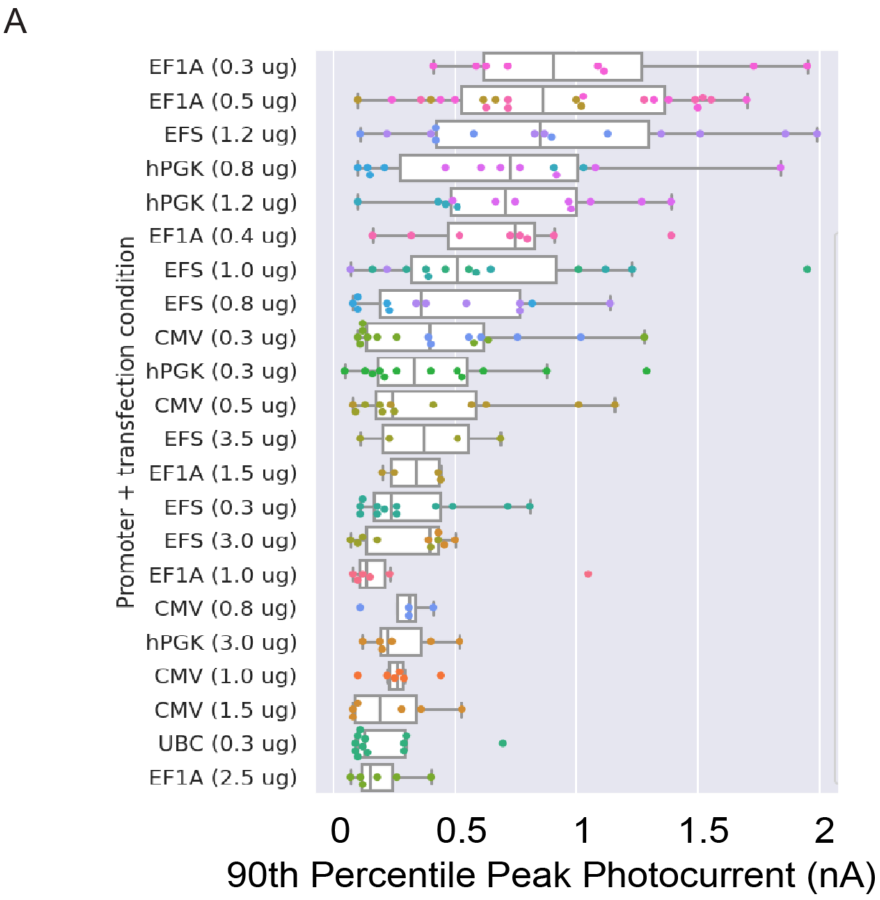


**Supplementary Figure 2.** Preliminary screen of promoters to drive expression of ChRmine
**(A)** Summary plot showing individual experiments (dots) and boxplots of autopatch photocurrent amplitude for HEK293T cells expressing ChRmine via lipofection. Y-axis shows the promoter and amount of DNA in micrograms per transfection used in the experiment.

### Supplementary Figure 3

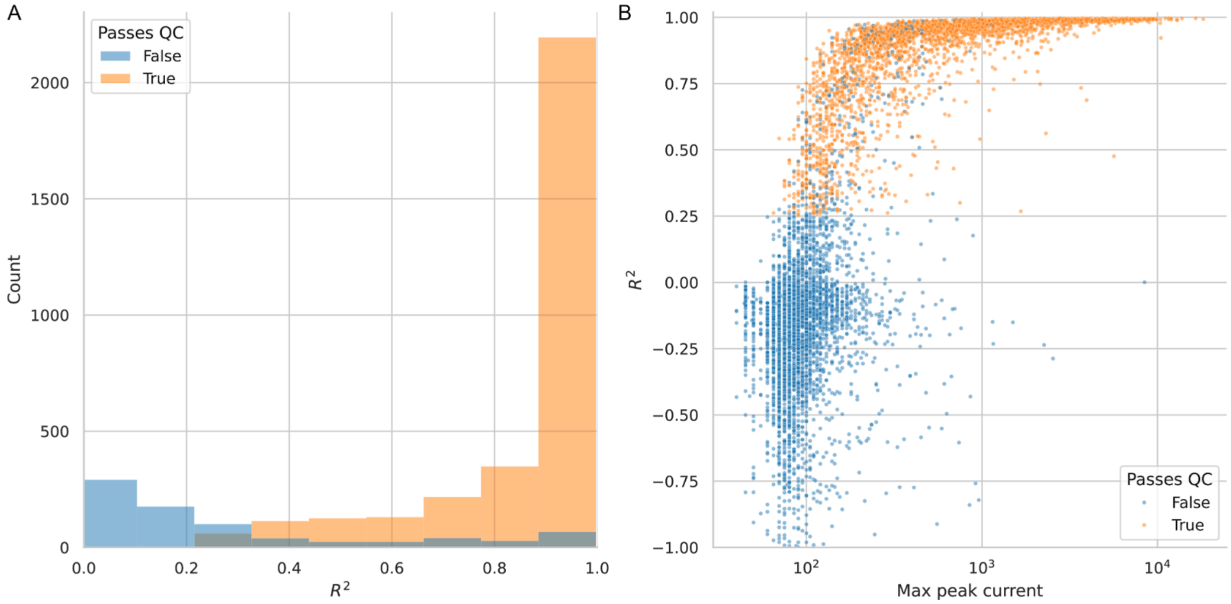


**Supplementary Figure 3.** Quality of model fits for individual autopatch recordings
**(A)** Median $R^2$ value was 0.96 for recordings where one stimulation condition evoked a median response of at least 200pA (passes QC, orange).
**(B)** Median $R^2$ improves with larger max peak current.

### Supplementary Figure 4

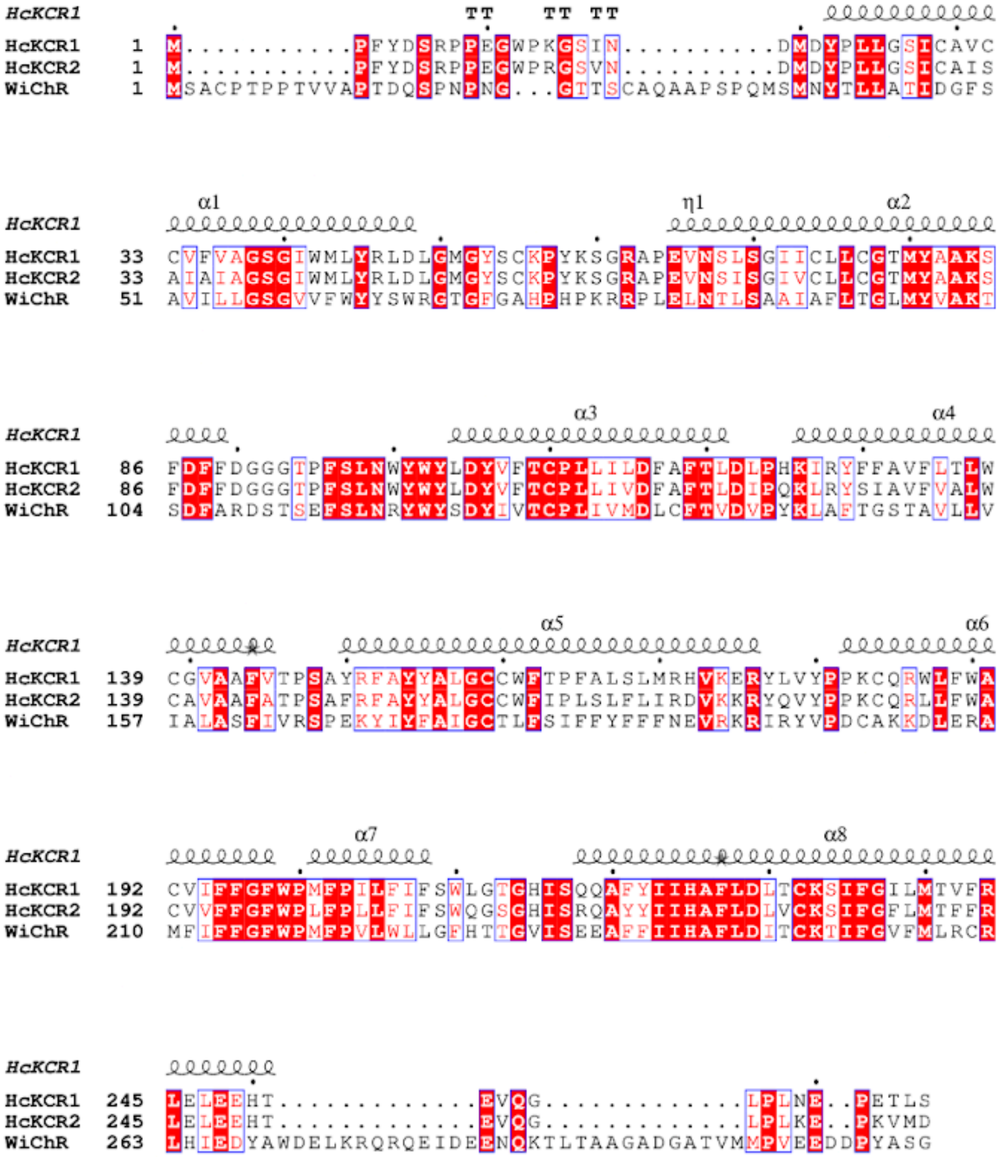


**Supplementary Figure 4.** Sequence alignment between HcKCR1, HcKCR2, and WiChR
**(A)** Biochemical studies of HcKCR1 and HcKCR2 have shown the Y222 residue to be involved in the $K^+$ selectivity filter. This corresponds to the F240 residue in the closely related WiChR (54% sequence similarity to HcKCR1).

### Supplementary Figure 5

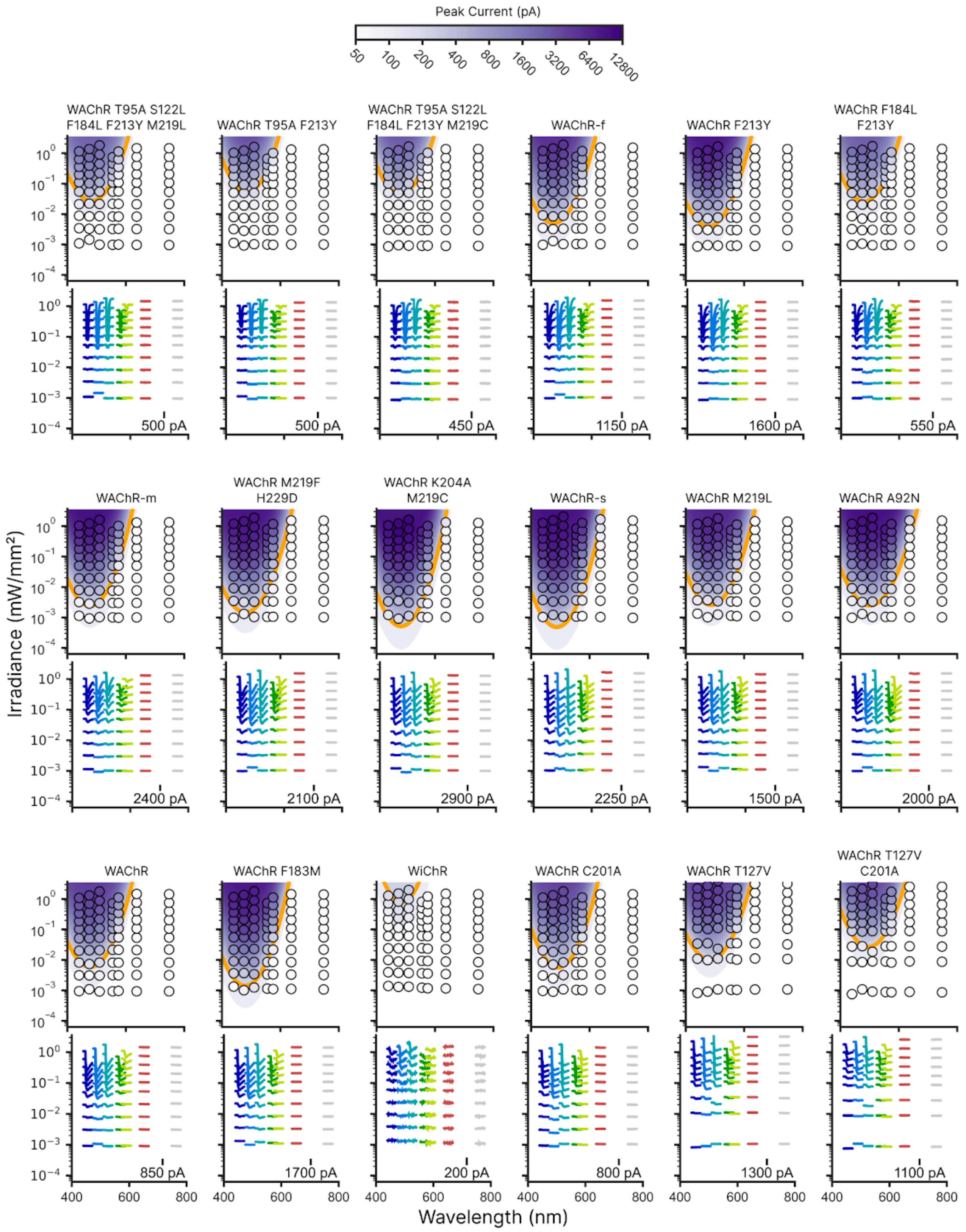


**Supplementary Figure 5.** Response surfaces and raw traces from some WAChR variants conducted using the autopatch assay during our screen.

### Supplementary Figure 6

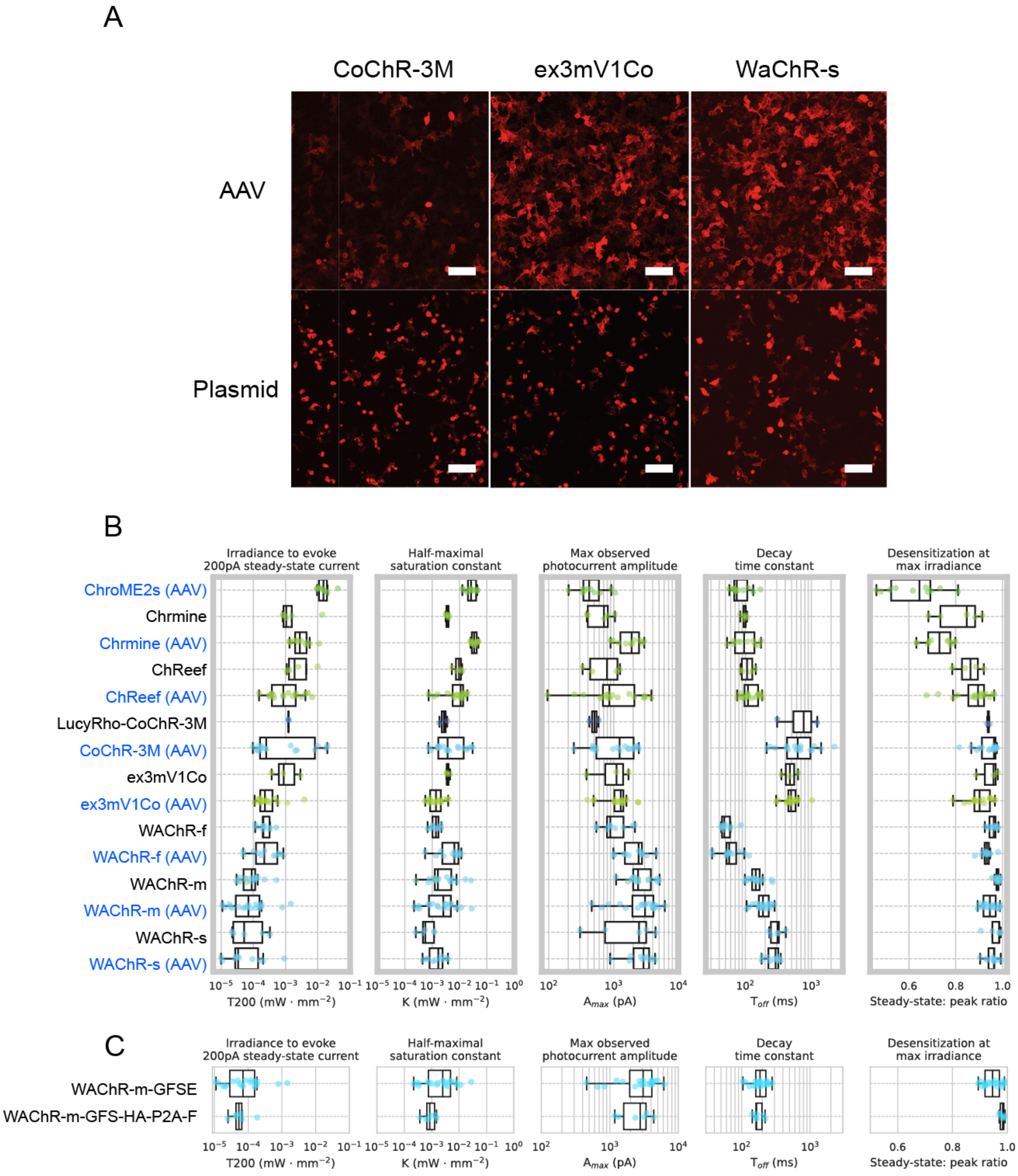


**Supplementary Figure 6.** Comparison of opsin expression and photocurrent from virus transduction and plasmid transfection **(A)** HEK293T cells expressing opsin with mScarlet tag by viral transduction with AAV (top row) or plasmid transfection (bottom row). Expression of CoChR-3M and ex3mV1Co with AAV results in better membrane localization and visual cell health compared to plasmid transfection. Plasmid transfection of WAChR-s maintains apparent cell health and membrane localization compared to virus transduction. WAChR-s plasmid expression appears healthier with better membrane localization compared to CoChR-3M and ex3mV1Co. Scale bars 50µm.

(**B**) Distributions of photocurrent metrics obtained from each opsin expressed either with AAV (blue text) or plasmid transfection (black text). Scatter point color indicates stimulation wavelength.
**(C)** Comparison of key performance metrics for the WAChR-m opsin as a fusion with mScarlet versus as a bicistronic construct. The first, WAChR-m-GFSE, represents the standard construct where the mScarlet fluorophore is fused to the opsin's C-terminus. The second, WAChR-m-GSE-HA-P2A-F, includes a P2A cleavage site, which separates the fluorophore from the opsin post-translation. Key functional properties, including activation threshold (T200), sensitivity (K), amplitude ($A_{max}$), speed ($\tau_{off}$), and desensitization are similar between the two constructs.

#### Supplementary Figure 7

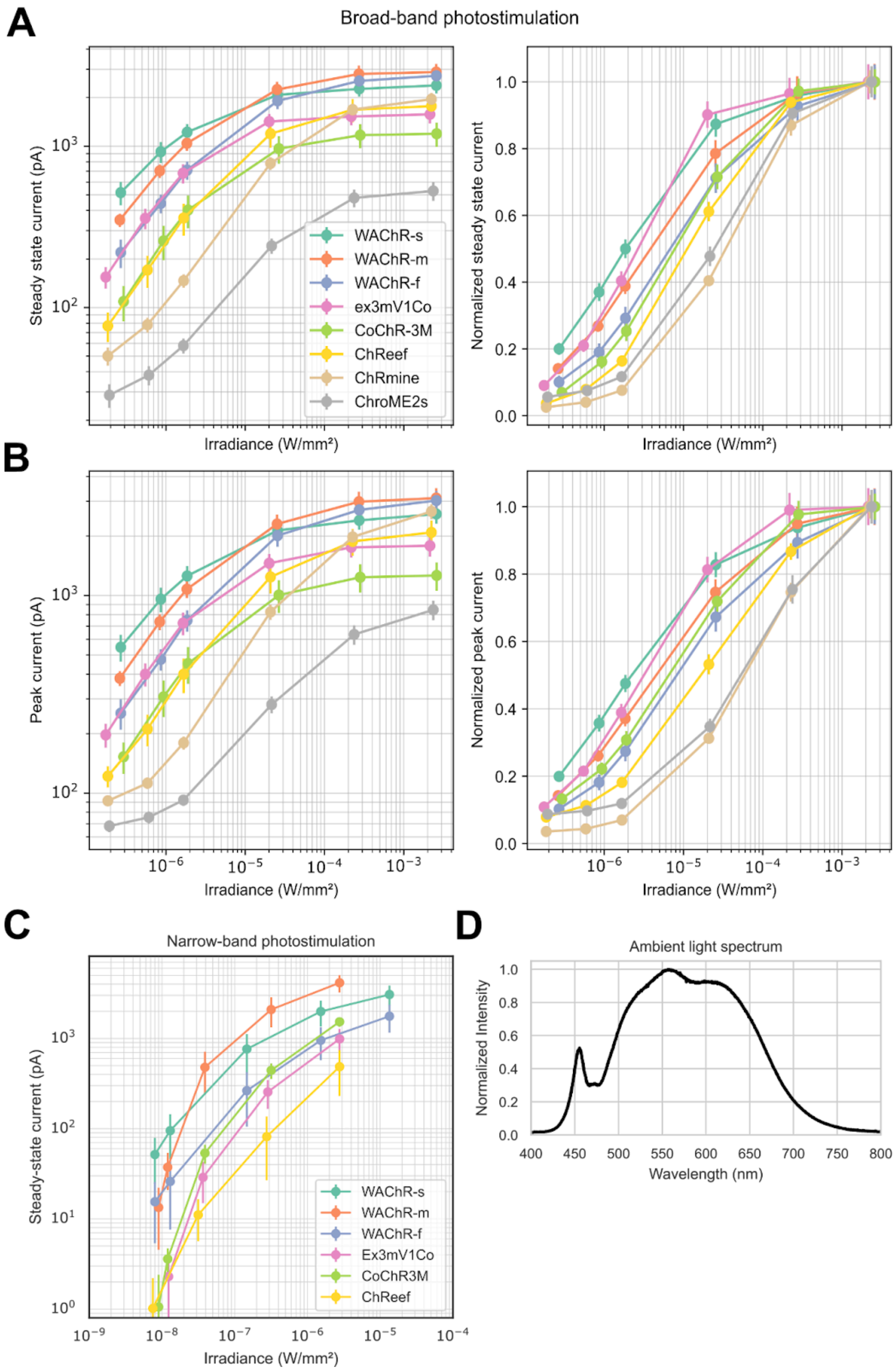


**Supplementary Figure 7.** Dose response relationships for WAChR variants and other sensitive opsins

**(A)** Dose response curves showing the steady-state photocurrents of various opsins in response to broad-band photostimulation across a range of irradiances. The left panel displays the absolute current in picoamperes (pA), while the right panel shows the same data normalized to the maximum response for each opsin.
**(B)** Dose response curves showing the peak photocurrents for the same opsins and conditions as in (A). The left panel displays the absolute peak current, and the right panel shows the normalized peak current.
**(C)** Steady-state current dose response curves for a subset of opsins measured using narrow-band photostimulation to probe sensitivity at low irradiances.
**(D)** Normalized spectrum from ambient indoor light as measured from the stage of the manual patch clamp rig.
All data were collected from manual whole-cell patch-clamp recordings of AAV-transduced HEK293T cells. Data points represent the mean, and error bars indicate the 95% confidence interval.

### Supplementary Figure 8

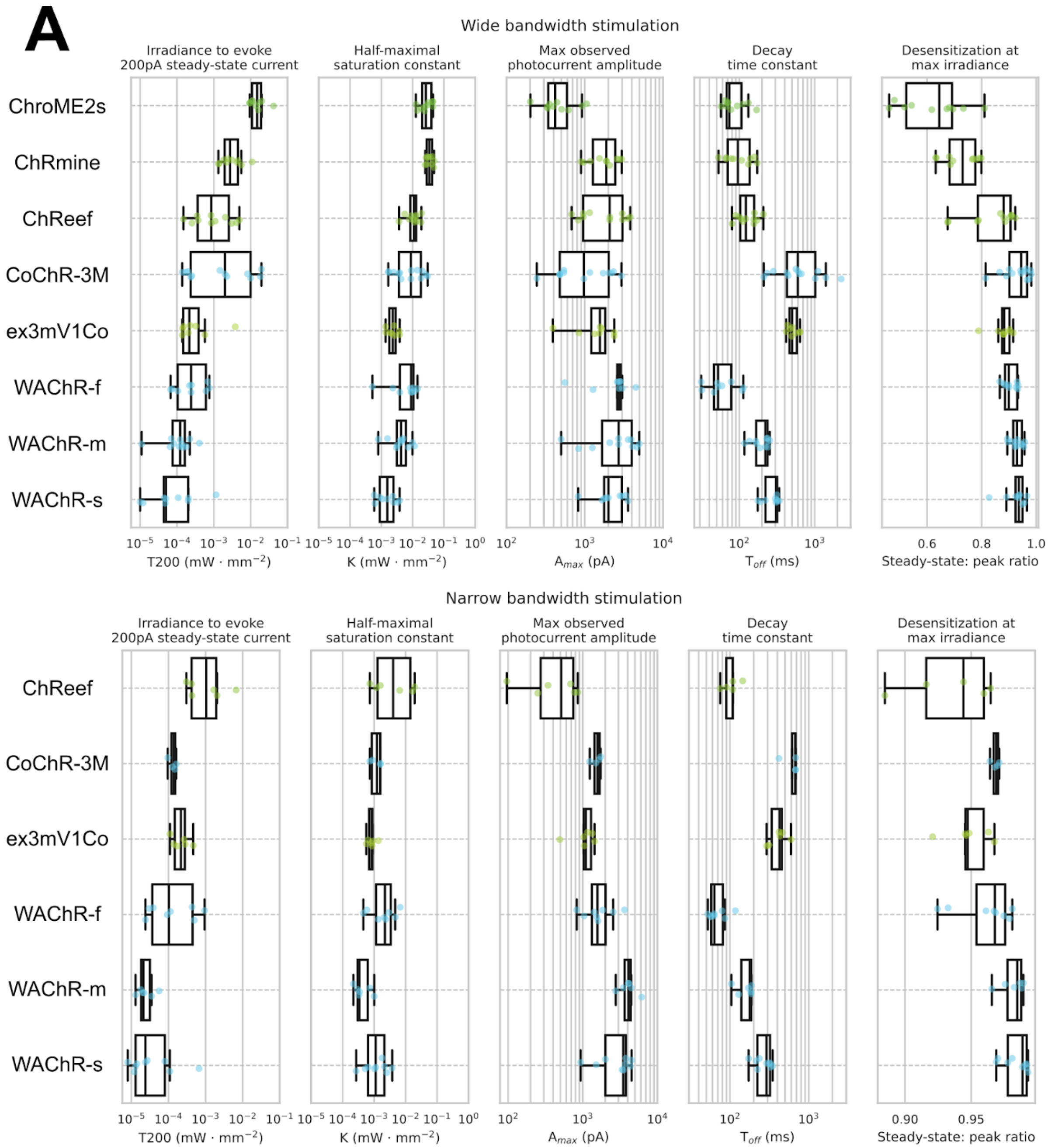


**Supplementary Figure 8**. Detailed sensitivity benchmarks of AAV-transduced opsins split by tunable light source recording configuration

(**A**) Distributions of photocurrent metrics obtained from fitting dose response curves to each opsin, as in Fig 5B, but broken down by experiments done in broad-band configuration (top) and narrow-band configuration (bottom)

### Supplementary Figure 9

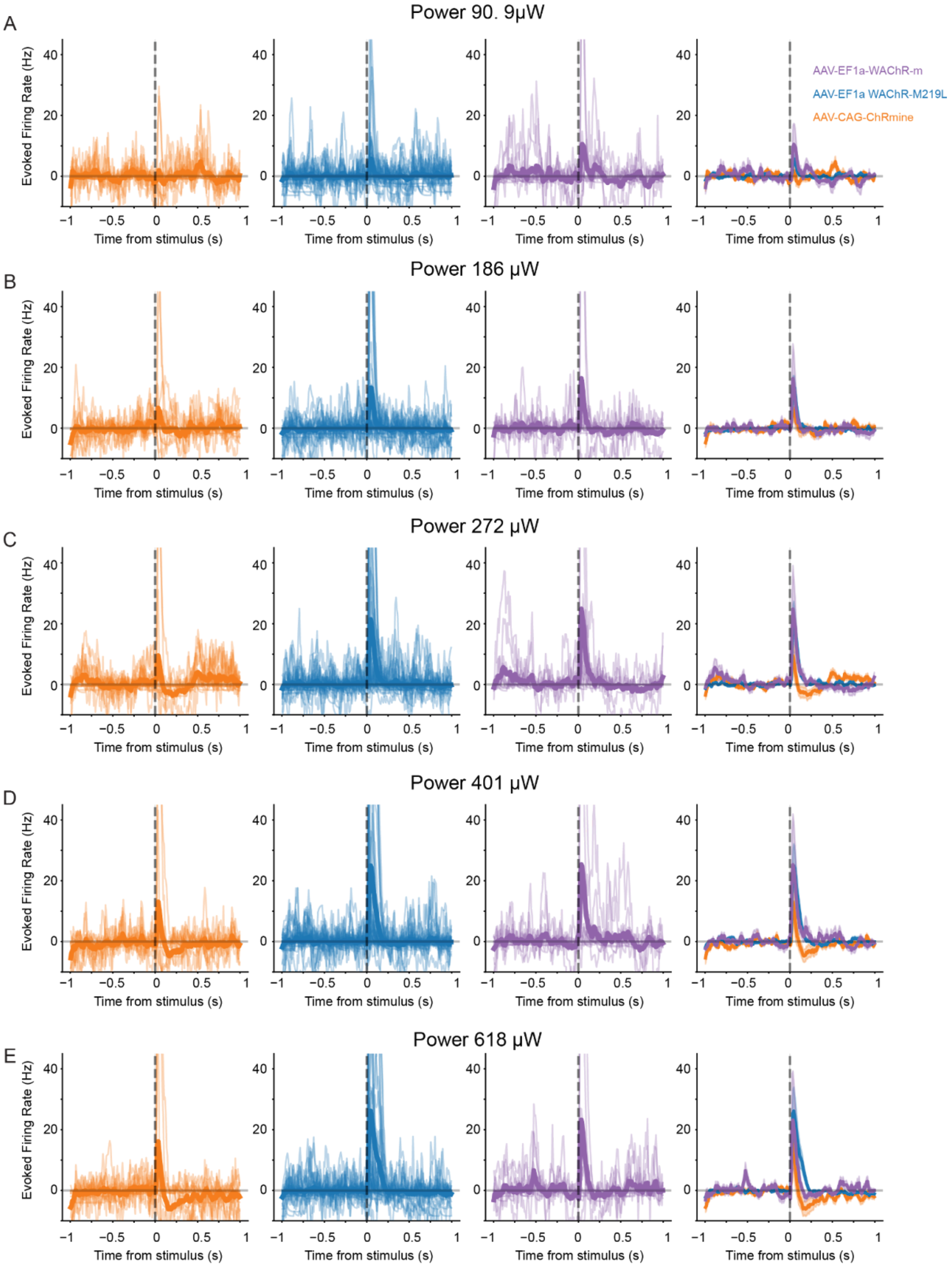

**Supplementary Figure 9.** Raw firing rates from *in vivo* electrophysiology recordings during 5 ms photostimulation
Mean firing rate (Hz) for units isolated from mice expressing AAV-CAG-ChRmine (orange), AAV-EF1a-WAChR-M219L (blue), or AAV-EF1a-WAChR-m (purple) during 5ms photostimulation trials with white light at power 90.9μW **(A)**, 186μW **(B)**, 272 μW **(C)**, 401 μW **(D)**, and 618 μW **(E)**.

### Supplementary Figure 10

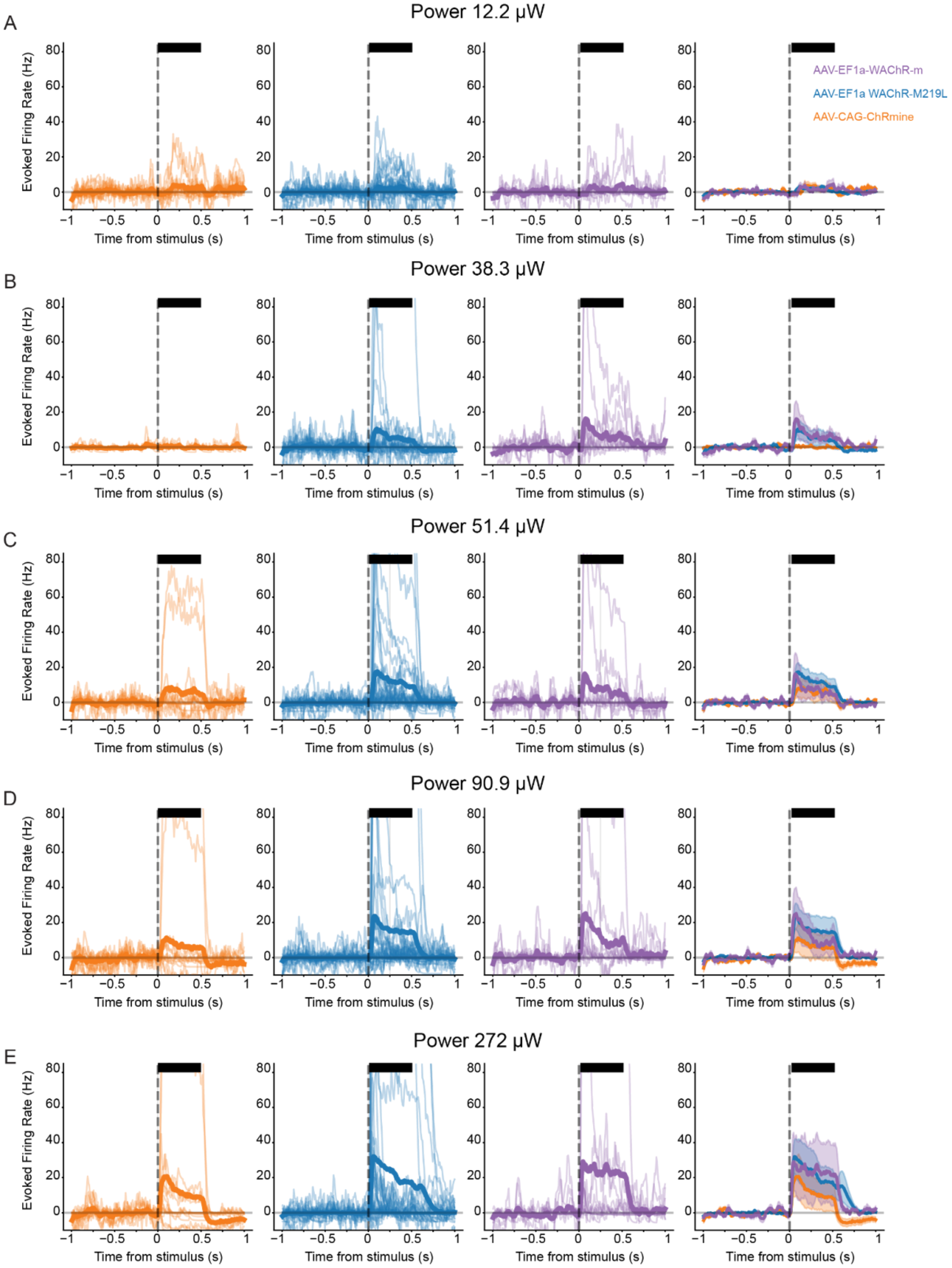


**Supplementary Figure 10.** Raw firing rates from *in vivo* electrophysiology recordings during 500ms photostimulation Mean firing rate (Hz) for units isolated from mice expressing AAV-CAG-ChRmine (orange), AAV-EF1a-WAChR-M219L (blue), or AAV-EF1a-WAChR-m (purple) during 500ms photostimulation trials with white light at power 12.2μW **(A)**, 38.3μW **(B)**, 51.4 μW **(C)**, 90.9 μW **(D)**, and 272 μW **(E)**.

### Supplementary Figure 11

A

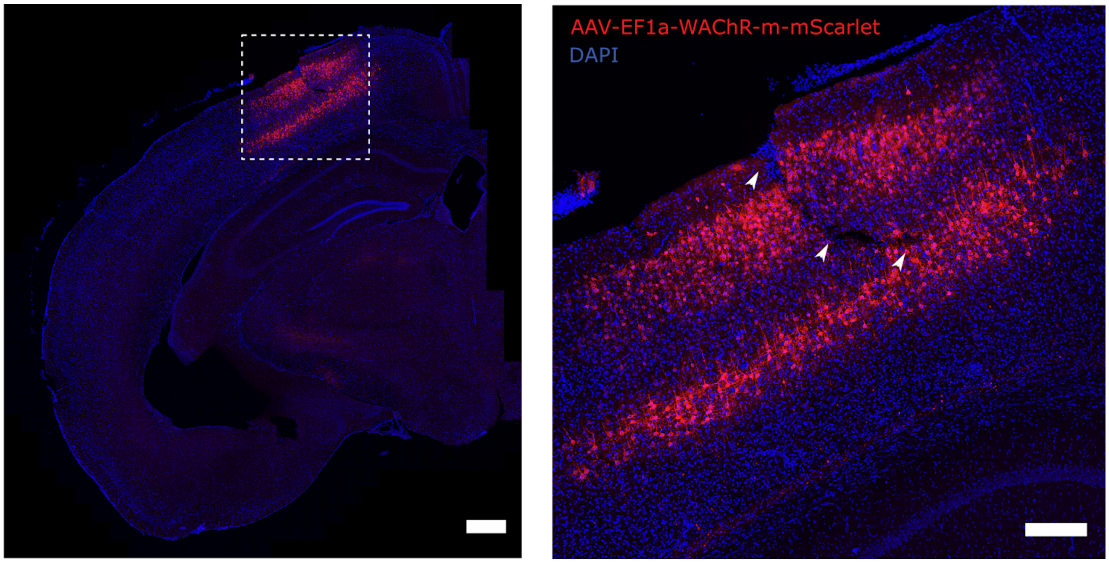


**Supplementary Figure 11.** Histology for AAV-EF1a-WAChR-M219L-mScarlet *in vivo* electrophysiology recording
**(A)** Confocal image of mouse cortex injected with AAV-EF1a-WAChR-M219L-mScarlet at location of neuropixels probe tract. Blue fluorescence is DAPI. Zoomed in image of inset on right (dashed lines). Arrows indicate neuropixels probe tract. Left scale bar 500 μm, right scale bar 200 μm.

### Supplementary Table 1

| Primer Name | Sequence |
|---|---|
| WiChR_F240A(pVBRP-EF1a)_F | CAAAAAAGCAGGCTGCCACCATGAGCGCCTGTCCTACACC |
| WiChR_F240A(pVBRP-EF1a)_R | ATCCTGCTCTTGGCGGCCGCTCCGCTAGCGTAGGGGTCGT |
| WiChR-A240_R | CCTCGCTGATCACGCCGGTGG |
| WiChR-A240-NRT_F | AGGCCTTTNRTATCATTCACG |
| WiChR-A240-MYG_F | AGGCCTTTMYGATCATTCACG |
| WiChR-A240-VAA_F | AGGCCTTTVAAATCATTCACG |
| WiChR-A240-DTT_F | AGGCCTTTDTTATCATTCACG |
| WiChR-A240-TGG_F | AGGCCTTTTGGATCATTCACG |
| WiChR-F239_R | CGCTGATCACGCCGGTGGTGT |
| WiChR-F239-VHG_F | AGGAGGCCVHGGCCATCATTC |
| WiChR-F239-BRT_F | AGGAGGCCBRTGCCATCATTC |
| WiChR-F239-ADC_F | AGGAGGCCADCGCCATCATTC |
| WiChR-F239-TGG_F | AGGAGGCCTGGGCCATCATTC |
| WiChR-W120_R | TCAGGCTAAACTCGCTGGTGC |
| WiChR-W120-VHG_F | ACAGGTACVHGTACAGCGACT |
| WiChR-W120-NRT_F | ACAGGTACNRTTACAGCGACT |
| WiChR-W120-WTT_F | ACAGGTACWTTTACAGCGACT |
| WiChR-M219_R | AGCCGAAGAAGATGAACATGG |
| WiChR-M219-NRT_F | TCTGGCCCNRTTTCCCCGTGC |
| WiChR-M219-VMG_F | TCTGGCCCVMGTTCCCCGTGC |
| WiChR-M219-DTT_F | TCTGGCCCDTTTTCCCCGTGC |
| WiChR-M219-TKG_F | TCTGGCCCTKGTTCCCCGTGC |
| WiChR-T127_R | AGTCGCTGTACCAGTACCTGT |
| WiChR-T127-NRT_F | ACATTGTGNRTTGCCCTCTGA |
| WiChR-T127-VWG_F | ACATTGTGVWGTGCCCTCTGA |
| WiChR-T127-SCG_F | ACATTGTGSCGTGCCCTCTGA |
| WiChR-T127-WTT_F | ACATTGTGWTTTGCCCTCTGA |
| WiChR-T127-TGG_F | ACATTGTGTGGTGCCCTCTGA |
| WiChR-C250_R | CTAGGAAGGCGTGAATGATGG |
| WiChR-C250-VHG_F | ACATTACCVHGAAGACCATTT |
| WiChR-C250-NAT_F | ACATTACCNATAAGACCATTT |
| WiChR-C250-VGC_F | ACATTACCVGCAAGACCATTT |
| WiChR-C250-WTT_F | ACATTACCWTTAAGACCATTT |
| WiChR-C250-TGG_F | ACATTACCTGGAAGACCATTT |

| | |
|---|---|
| WiChR-T252_R | TAATGTCTAGGAAGGCGTGAA |
| WiChR-T252-NRT_F | CCTGCAAGNRTATTTTCGGCG |
| WiChR-T252-VWG_F | CCTGCAAGVWGATTTTCGGCG |
| WiChR-T252-SCG_F | CCTGCAAGSCGATTTTCGGCG |
| WiChR-T252-WTT_F | CCTGCAAGWTTATTTTCGGCG |
| WiChR-T252-TGG_F | CCTGCAAGTGGATTTTCGGCG |
| WiChR-I131_R | AGGTCACAATGTAGTCGCTGT |
| WiChR-I131-VHG_F | GCCCTCTGVHGGTGATGGACT |
| WiChR-I131-NRT_F | GCCCTCTGNRTGTGATGGACT |
| WiChR-I131-TTT_F | GCCCTCTGTTTGTGATGGACT |
| WiChR-I131-TGG_F | GCCCTCTGTGGGTGATGGACT |

**Supplementary Table 1.** Primer sequences used in mutagenesis

### Supplementary Table 2

| Name | Sequence |
| --- | --- |
| pEF1A-WAChR-GFSE<br>(Underline = EF1a, Bold = WAChR, Italic = mScarlet) | CAACTTTGTATAGAAAAGTTGGGCTCCGGTGCCCGTCAGTGGGCAGAGCGCACATCGC<br>CCACAGTCCCCGAGAAGTTGGGGGGAGGGGTCGGCAATTGAACCGGTGCCTAGAGAAG<br>GTGGCGCGGGGTAAACTGGGAAAGTGATGTCGTGTACTGGCTCCGCCTTTTTCCCGAG<br>GGTGGGGGAGAACCGTATATAAGTGCAGTAGTCGCCGTGAACGTTCTTTTTCGCAACG<br>GGTTTGCCGCCAGAACACAGGTAAGTGCCGTGTGTGGTTCCCGCGGGCCTGGCCTCTT<br>TACGGGTTATGGCCCTTGCGTGCCTTGAATTACTTCCACCTGGCTGCAGTACGTGATT<br>CTTGATCCCGAGCTTCGGGTTGGAAGTGGGTGGGAGAGTTCGAGGCCTTGCGCTTAAG<br>GAGCCCCTTCGCCTCGTGCTTGAGTTGAGGCCTGGCCTGGGCGCTGGGGCCGCCGCGT<br>GCGAATCTGGTGGCACCTTCGCGCCTGTCTCGCTGCTTTCGATAAGTCTCTAGCCATT<br>TAAAATTTTTGATGACCTGCTGCGACGCTTTTTTTCTGGCAAGATAGTCTTGTAAATG<br>CGGGCCAAGATCTGCACACTGGTATTTCGGTTTTTGGGGCCGCGGGCGGCGACGGGGC<br>CCGTGCGTCCCAGCGCACATGTTCGGCGAGGCGGGGCCTGCGAGCGCGGCCACCGAGA<br>ATCGGACGGGGGTAGTCTCAAGCTGGCCGGCCTGCTCTGGTGCCTGGTCTCGCGCCGC<br>CGTGTATCGCCCCGCCCTGGGCGGCAAGGCTGGCCCGGTCGGCACCAGTTGCGTGAGC<br>GGAAAGATGGCCGCTTCCCGGCCCTGCTGCAGGGAGCTCAAAATGGAGGACGCGGCGC<br>TCGGGAGAGCGGGCGGGTGAGTCACCCACACAAAGGAAAAGGGCCTTTCCGTCCTCAG<br>CCGTCGCTTCATGTGACTCCACGGAGTACCGGGCGCCGTCCAGGCACCTCGATTAGTT<br>CTCGAGCTTTTGGAGTACGTCGTCTTTAGGTTGGGGGGAGGGGTTTTATGCGATGGAG<br>TTTCCCCACACTGAGTGGGTGGAGACTGAAGTTAGGCCAGCTTGGCACTTGATGTAAT<br>TCTCCTTGGAATTTGCCCTTTTTGAGTTTGGATCTTGGTTCATTCTCAAGCCTCAGAC<br>AGTGGTTCAAAGTTTTTTTCTTCCATTTCAGGTGTCGTGACAAGTTTGTACAAAAAAG<br>CAGGCTGCCACC**ATGAGCGCCTGTCCTACACCCCCTACCGTGGTGGCCCCCACAGATC**<br>**AAAGCCCCAATCCTAACGGAGGCACCACCTCTTGTGCTCAGGCTGCCCCTTCTCCTCA**<br>**GATGAGCATGAATTACACCCTGCTTGCAACCATCGACGGATTTAGCGCCGTGATTCTG**<br>**CTTGGAAGCGGCGTGGTGTTCTGGTACTACTCTTGGAGGGGAACCGGCTTTGGAGCCC**<br>**ACCCCCATCCCAAGAGAAGACCCCTGGAACTGAATACCCTGTCTGCCGCCATCGCCTT**<br>**CTTAACCGGCCTGATGTACGTGGCCAAGACCAGCGACTTTGCTAGGGACAGCACCAGC**<br>**GAGTTTAGCCTGAACAGGTACTGGTACAGCGACTACATTGTGACCTGCCCTCTGATTG**<br>**TGATGGACTTATGTTTCACCGTTGACGTTCCCTACAAGCTGGCATTCACCGGAAGCAC**<br>**CGCCGTGCTGCTGGTGATTGCCTTAGCCAGCTTTATTGTGAGGAGCCCCGAGAAGTAT**<br>**ATTTACTTTGCAATTGGCTGCACACTGTTTAGCATTTTCTTTTACTTCTTTTTCAATG**<br>**AGGTGAGAAAGAGGATTAGGTACGTGCCCGACTGCGCCAAGAAGGACCTGGAGAGGGC**<br>**CATGTTCATCTTCTTCGGCTTCTGGCCCATGTTCCCCGTGCTGTGGCTGCTGGGCTTT**<br>**CACACCACCGGCGTGATCAGCGAGGAGGCCTTTGCTATCATTCACGCCTTCCTAGACA**<br>**TTACCTGCAAGACCATTTTCGGCGTGTTCATGCTGAGGTGCAGACTGCACATCGAGGA**<br>**CTACGCCTGGGATGAGCTGAAAAGGCAGAGACAGGAGATCGACGAGGAGAATCAGAAA**<br>**ACCTTGACCGCCGCTGGCGCCGATGGCGCCACCGTGATGATGCCCGTTGAAGAAGACG**<br>**ACCCCTACGCTAGCGGA**GCGGCCGCCAAGAGCAGGATCACCAGCGAGGGCGAGTACAT<br>CCCCCTGGACCAGATCGATATCAACGTGG*GTGAGCAAGGGCGAGGCAGTGATCAAGGAGT*<br>*TCATGCGGTTCAAGGTGCACATGGAGGGCTCCATGAACGGCCACGAGTTCGAGATCGAGG*<br>*GCGAGGGCGAGGGCCGCCCCTACGAGGGCACCCAGACCGCCAAGCTGAAGGTGACCAAGG*<br>*GTGGCCCCCTGCCCTTCTCCTGGGACATCCTGTCCCCTCAGTTCATGTACGGCTCCAGGG*<br>*CCTTCACCAAGCACCCCGCCGACATCCCCGACTACTATAAGCAGTCCTTCCCCGAGGGCT*<br>*TCAAGTGGGACCGCGTGATGAACTTCGAGGACGGCGGCGCCGTGACCGTGACCCAGGACA*<br>*CCTCCCTGGAGGACGGCACCCTGATCTACAAGGTGAAGCTCCGCGGCACCAACTTCCCTC*<br>*CTGACGGCCCCGTAATGCAGAAGAAGACAATGGGCTGGGAAGCGTCCACCGAGCGGTTGT*<br>*ACCCCGAGGACGGCGTGCTGAAGGGCGACATTAAGATGGCCCTGCGCCTGAAGGACGGCG*<br>*GGCGCTACCTGGCGGACTTCAAGACCACCTACAAGGCCAAGAAGCCCGTGCAGATGCCCG*<br>*GCGCCTACAACGTGGACCGCAAGTTGGACATCACCTCCCACAACGAGGACTACACCGTGG*<br>*TGGAACAGTACGAACGCTCCGAGGGCCGCCACTCCACCGGCGGCATGGACGAGCTGTACA*<br>*AG*CAGTCCCAGCCCATCCTCAACACCAAGGAGATGGCCCCGCAGAGCAAGCCTCCAGA<br>GGAGCTGGAGATGAGCAGCATGCCCAGCCCCGTGGCCCCTCTGCCCGCACGCACGGAG<br>GGCGTCATCGACATGCGGAGCATGTCCAGCATTGACAGCTTCATCAGCTGTGCCACGG<br>ACTTCCCTGAAGCCACCAGATTCTTCTGCTACGAGAACGAGGTGTAAACCCAGCTTTC<br>TTGTACAAAGTGGTGATGGCCGGCCGCTTCGAGCAGACATGATAAGATACATTGATGA |

| | |
|---|---|
| | GTTTGGACAAACCACAACTAGAATGCAGTGAAAAAAATGCTTTATTTGTGAAATTTGT<br>GATGCTATTGCTTTATTTGTAACCATTATAAGCTGCAATAAACAAGTTAACAACAACA<br>ATTGCATTCATTTTATGTTTCAGGTTCAGGGGGAGGTGTGGGAGGTTTTTTAAAGCAA<br>GTAAAACCTCTACAAATGTGGTAGCGGCCGCGGCGCTCTTCCGCTTCCTCGCTCACTG<br>ACTCGCTGCGCTCGGTCGTTCGGCTGCGGCGAGCGGTATCAGCTCACTCAAAGGCGGT<br>AATACGGTTATCCACAGAATCAGGGGATAACGCAGGAAAGAACATGTGAGCAAAAGGC<br>CAGCAAAAGGCCAGGAACCGTAAAAAGGCCGCGTTGCTGGCGTTTTTCCATAGGCTCC<br>GCCCCCCTGACGAGCATCACAAAAATCGACGCTCAAGTCAGAGGTGGCGAAACCCGAC<br>AGGACTATAAAGATACCAGGCGTTTCCCCCTGGAAGCTCCCTCGTGCGCTCTCCTGTT<br>CCGACCCTGCCGCTTACCGGATACCTGTCCGCCTTTCTCTCTTCGGGAAGCGTGGCGC<br>TTTCTCATAGCTCACGCTGTAGGTATCTCAGTTCGGTGTAGGTCGTTCGCTCCAAGCT<br>GGGCTGTGTGCACGAACCCCCCGTTCAGCCCGACCGCTGCGCCTTATCCGGTAACTAT<br>CGTCTTGAGTCCAACCCGGTAAGACACGACTTATCGCCACTGGCAGCAGCCACTGGTA<br>ACAGGATTAGCAGAGCGAGGTATGTAGGCGGTGCTACAGAGTTCTTGAAGTGGTGGCC<br>TAACTACGGCTACACTAGAAGAACAGTATTTGGTATCTGCGCTCTGCTGAAGCCAGTT<br>ACCTTCGGAAAAAGAGTTGGTAGCTCTTGATCCGGCAAACAAACCACCGCTGGTAGCG<br>GTGGTTTTTTTGTTTGCAAGCAGCAGATTACGCGCAGAAAAAAAGGATCTCAAGAAGA<br>TCCTTTGATCTTTTCTACGGGGTCTGACGCTCAGTGGAACGAAAACTCACGTTAAGGG<br>ATTTTGGTCATGAGATTATCAAAAAGGATCTTCACCTAGATCCTTTTAAATTAAAAAT<br>GAAGTTTTAAATCAATCTAAAGTATATATGAGTAAACTTGGTCTGACAGTTACCAATG<br>CTTAATCAGTGAGGCACCTATCTCAGCGATCTGTCTATTTCGTTCATCCATAGTTGCC<br>TGACTCCCCGTCGTGTAGATAACTACGATACGGGAGGGCTTACCATCTGGCCCCAGTG<br>CTGCAATGATACCGCGAGACCCACGCTCACCGGCTCCAGATTTATCAGCAATAAACCA<br>GCCAGCCGGAAGGGCCGAGCGCAGAAGTGGTCCTGCAACTTTATCCGCCTCCATCCAG<br>TCTATTAATTGTTGCCGGGAAGCTAGAGTAAGTAGTTCGCCAGTTAATAGTTTGCGCA<br>ACGTTGTTGCCATTGCTACAGGCATCGTGGTGTCACGCTCGTCGTTTGGTATGGCTTC<br>ATTCAGCTCCGGTTCCCAACGATCAAGGCGAGTTACATGATCCCCCATGTTGTGCAAA<br>AAAGCGGTTAGCTCCTTCGGTCCTCCGATCGTTGTCAGAAGTAAGTTGGCCGCAGTGT<br>TATCACTCATGGTTATGGCAGCACTGCATAATTCTCTTACTGTCATGCCATCCGTAAG<br>ATGCTTTTCTGTGACTGGTGAGTACTCAACCAAGTCATTCTGAGAATAGTGTATGCGG<br>CGACCGAGTTGCTCTTGCCCGGCGTCAATACGGGATAATACCGCGCCACATAGCAGAA<br>CTTTAAAAGTGCTCATCATTGGAAAACGTTCTTCGGGGCGAAAACTCTCAAGGATCTT<br>ACCGCTGTTGAGATCCAGTTCGATGTAACCCACTCGTGCACCCAACTGATCTTCAGCA<br>TCTTTTACTTTCACCAGCGTTTCTGGGTGAGCAAAAACAGGAAGGCAAAATGCCGCAA<br>AAAAGGGAATAAGGGCGACACGGAAATGTTGAATACTCATACTCTTCCTTTTTCAATA<br>TTATTGAAGCATTTATCAGGGTTATTGTCTCATGAGCGGATACATATTTGAATGTATT<br>TAGAAAAATAAACAAATAGGGGTTCCGCGCACATTTCCCCGAAAAGTGCCACCTGACG<br>TCTAAGAAACCATTATTATCATGACATTAACCTATAAAAATAGGCGTATCACGAGGCC<br>CTTTCGTCGGCGCGCCGCGGCCGC |

**Supplementary Table 2.** Plasmid Sequences
